# The pAblo·pCasso self-curing vector toolset for unconstrained cytidine and adenine base-editing in Gram-negative bacteria

**DOI:** 10.1101/2023.04.16.537106

**Authors:** Ekaterina Kozaeva, Zacharias S. Nielsen, Manuel Nieto-Domínguez, Pablo I. Nikel

## Abstract

A synthetic biology toolkit, exploiting clustered regularly interspaced short palindromic repeats (CRISPR) and modified CRISPR-associated protein (Cas) base-editors, was developed for genome engineering in Gram-negative bacteria. Both a cytidine base-editor (CBE) and an adenine base-editor (ABE) have been optimized for precise single-nucleotide modification of plasmid and genome targets. CBE comprises a cytidine deaminase conjugated to a Cas9 nickase from *Streptococcus pyogenes* (*^Sp^*nCas9), resulting in C→T (or G→A) substitutions. Conversely, ABE consists of an adenine deaminase fused to *^Sp^*nCas9 for A→G (or T→C) editing. Several nucleotide substitutions were achieved using these plasmid-borne base-editing systems and a novel protospacer adjacent motif (PAM)-relaxed *^Sp^*nCas9 (SpRY) variant. Base-editing was validated in *Pseudomonas putida* and other Gram-negative bacteria by inserting premature *STOP* codons into target genes, thereby inactivating both fluorescent proteins and metabolic (antibiotic-resistance) functions. The functional knockouts obtained by engineering *STOP* codons *via* CBE were reverted to the wild-type genotype using ABE. Additionally, a series of induction-responsive vectors was developed to facilitate the curing of the base-editing platform in a single cultivation step, simplifying complex strain engineering programs without relying on homologous recombination and yielding plasmid-free, modified bacterial cells.

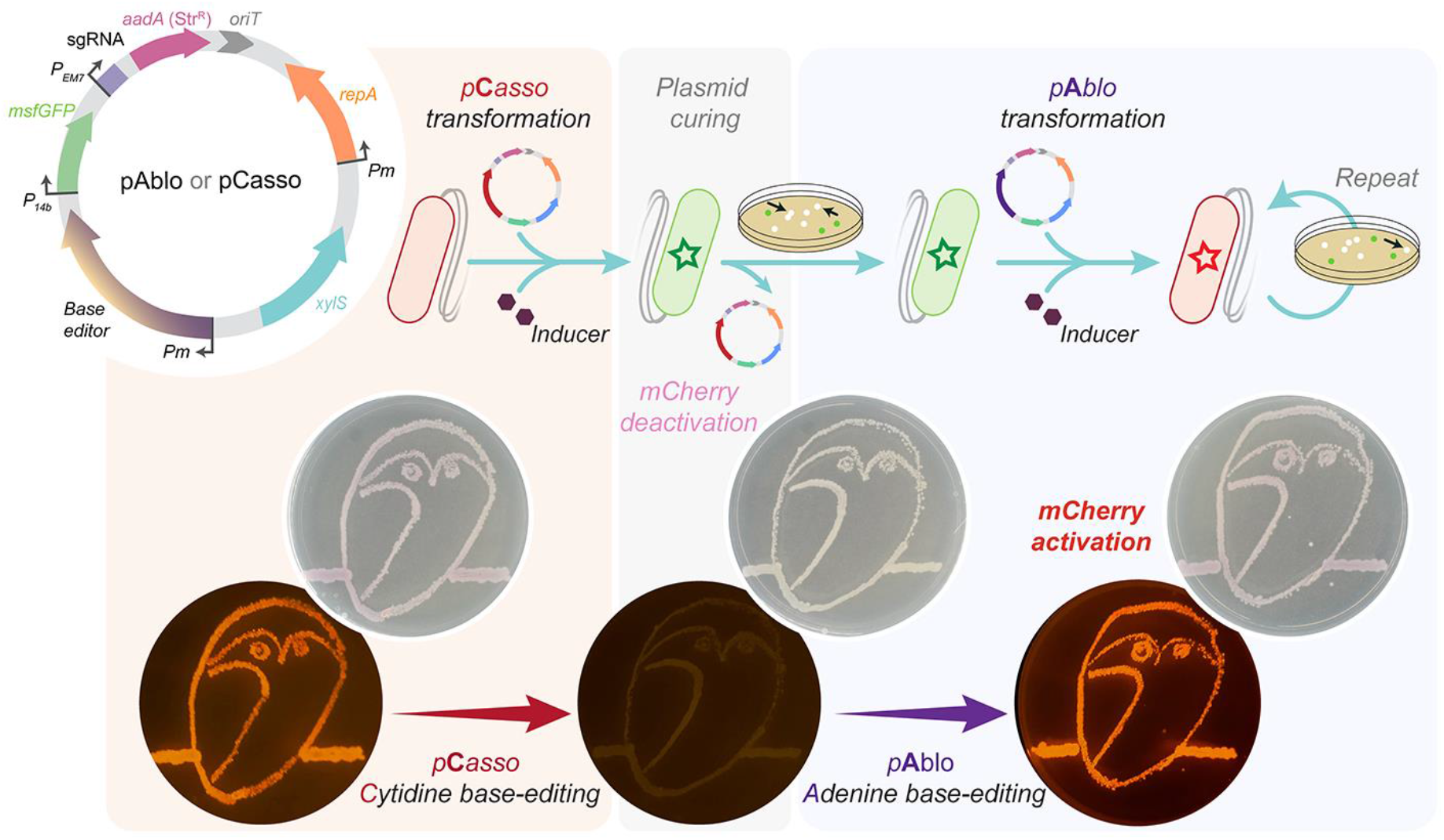

## INTRODUCTION

Precision genome editing continues to revolutionize life sciences, with a direct impact in multiple fields (Wang *et al*., 2022; Wang and Doudna, 2023)—including both fundamental studies and applied sciences, e.g. metabolic engineering (Cho *et al*., 2018; Zhao *et al*., 2021; Volke *et al*., 2023). Among these emerging approaches, base-editing of bacterial genomes assisted by clustered regularly interspaced short palindromic repeats (CRISPR) and their associated proteins (Cas), allows for single-nucleotide-resolution modification of genomic sites in a homologous recombination-independent fashion (Anzalone *et al*., 2020; Nishida and Kondo, 2020; Sun *et al*., 2020; Walton *et al*., 2020). Such DNA engineering techniques rely on synthetic editing modules—most commonly, cytidine base-editors (CBE) that result in a cytidine (C)→thymidine (T) [or guanine (G)→adenine (A)] substitution, and adenine base-editors (ABE) that enable A→G (or T→C) modifications (Komor *et al*., 2016). A CBE typically carries a cytidine deaminase fused to a Cas9 nickase (nCas9) and uracil glycosylase inhibitors (Komor *et al*., 2017), and an ABE consists of an nCas9 and an evolved adenine deaminase, capable of recognizing single-stranded DNA (ssDNA) (Gaudelli *et al*., 2017). In the type-II CRISPR-Cas system, the naturally-occurring CRISPR-associated protein 9 from *Streptococcus pyogenes* (*^Sp^*Cas9) recognizes its targets with the help of the cognate guide, a complex of a transactivating crRNA (tracrRNA) and a CRISPR RNA (crRNA) transcribed from the CRISPR array (Sapranauskas *et al*., 2011). As a defense system from invading nucleic acid fragments, a CRISPR array functions as a library of spacers, adapted from DNA structures located next to specific short sequences called protospacer-adjacent motifs (PAMs). A PAM is required for binding of the Cas9 domain to the target sequence (Jiang *et al*., 2013). In systems engineered for base-editing, the nCas9 nickase (displaying a D10A mutation that deactivates the RuvC cleavage domain of *^Sp^*Cas9) catalyzes a single-strand DNA break upon binding to a compatible PAM, relying on the active HNH cleavage domain. The single-strand break is introduced in the target DNA strand through the recognition of a specific synthetic single-guide RNA (sgRNA), and the cleavage is constrained by the identity of the PAM sequence (Gasiunas *et al*., 2012). *^Sp^*Cas9 recognizes the 5’-*N*GG-3’ motif, where *N* can be any nucleotide. From a practical point of view, however, the PAM recognition profile may constrain CRISPR-mediated genome engineering, leaving numerous loci inaccessible to editing—e.g. regions with low G+C content. To tackle this issue, Cas proteins have been optimized for flexible PAM recognition (Walton *et al*., 2020). These engineered variants hold great potential for improved genome editing as they render more DNA sites accessible to several types of modifications. Furthermore, fusing such proteins with nucleotide deaminase domains provides a unique opportunity for modification of almost any DNA base of choice without narrow PAM recognition. Developing this type of base-editing methodology is especially relevant for microbial strain engineering, as it would allow for fast genome modifications along the entire bacterial chromosome. PAM-independent base-editing by CBEs enables the integration of premature *STOP* codons into an open reading frame (ORF), such that the 5’-CAG-3’, 5’-CAA-3’ or 5’-CGA-3’ codons are converted into 5’-TAG-3’ (amber), 5’-TAA-3’ (ochre) or 5’-TGA-3’ (opal) triplets, respectively—thereby inactivating the cognate protein (i.e. functional knockout). These modifications could be reverted to restore the original (coding) sequence of the target ORFs using ABEs, enabling a functional switch for genome modifications. Although some efforts have been made to partially develop these technologies in model bacteria, e.g. *Escherichia coli* (Schultenkämper *et al*., 2020; Shelake *et al*., 2022), the availability of base-editing tools for alternative microbial hosts (e.g. *Pseudomonas* species) is restricted to a few exploratory examples.

The *Pseudomonas* genus comprises key microbial models for both fundamental and applied research. *Pseudomonas* species are known to tolerate high levels of abiotic stresses, e.g. oxidative damage and solvents, underscoring their potential as hosts for metabolic engineering. For instance, *P. putida* displays a versatile metabolism that makes this species an ideal platform for biotechnology (Volke *et al*., 2020a; Weimer *et al*., 2020). Clinically-relevant species, e.g. *P. aeruginosa*, are a frequent cause of nosocomial complications in immunosuppressed patients, whereas other *Pseudomonas* are phytopathogens that can infect a variety of commercially-relevant crops (Brinkman *et al*., 2021). The advent of genome engineering tools has substantially propelled research on *Pseudomonas*, partly elucidating complex connections between genotypes and phenotypes (Volke and Nikel, 2018; Kozaeva *et al*., 2021). Tools for gene deletion, insertion and genome modification(s) are well established for *P. putida* (Martínez-García and de Lorenzo, 2019). However, these methods often prove labor-intensive, demanding and difficult to automate. Also, non-homologous end joining (NHEJ) mechanisms are virtually absent in prokaryotes (Öz *et al*., 2021), hindering the introduction of frameshifts in target ORFs and subsequent functional knock-outs. A few recent studies that employed CRISPR-Cas gene editing in *Pseudomonas* species have showcased the feasibility and precision of this approach in genome engineering (Abdullah *et al*., 2022; Volke *et al*., 2022; Yue *et al*., 2022).

In this work, we describe a novel application of an engineered, nearly-PAM-less *^Sp^*Cas9 variant (SpRY) for precise nucleotide substitution in Gram-negative bacteria by developing a plasmid toolset (termed *pAblo·pCasso*) for efficient base-editing. Different types of editor modules (i.e. CBE and ABE) were characterized using both the canonical *^Sp^*nCas9 and the nearly-PAM-less SpRY proteins. SpRY can target almost all PAMs in the genome (with a 5’-*N*R*N*-3’ motif preferred over 5’-*N*Y*N*-3’, where R is A or G and Y is C or T) (Walton *et al*., 2020). Furthermore, we developed a set of plasmids that are induction-dependent and self-curing, featuring conditional replication to simplify vector elimination. These vectors were exploited as a platform for precision base-editing in three different bacterial species for targets located both in the chromosome and in a separate plasmid. Hence, the genome editing techniques established in this study, together with streamlined protocols for vector curing, facilitate genome engineering in Gram-negative bacteria, with a significant potential for expediting research in non-model microbial species.

## MATERIALS AND METHODS

### Bacterial strains, plasmids and culture conditions

Bacterial strains and plasmids used in this study are listed in **Table 1**. *E. coli*, *P. putida* and *P. fluorescens* cultures were grown in lysogeny broth (LB) medium (10 g L^−1^ tryptone, 5 g L^−1^ yeast extract and 10 g L^−1^ NaCl; solid culture media additionally contained 15 g L^−1^ agar) (Nikel *et al*., 2008) at 37°C (*E. coli*) and 30°C (*P. putida* and *P. fluorescens*) (Nikel *et al*., 2010; Green and Sambrook, 2012). Streptomycin (Str) was added whenever needed at 100 μg mL^−1^ (for *E. coli* cultures) or 200 μg mL^−1^ (for *P. putida* cultures); kanamycin (Km) was used at 50 μg mL^−1^. In experiments involving base-editing of the *nicX* gene, nicotinic acid was added to culture media at 5 mM. Liquid cultures were agitated at 200 rpm (MaxQ™ 8000 incubator; ThermoFisher Scientific, Waltham, MA, USA). The optical density measured at 600 nm (OD_600_) was recorded in a Genesys 20 spectrophotometer (ThermoFisher Scientific) to estimate cell density. For preculture preparation, four independent colonies were picked from an LB medium plate and used to inoculate 96-deep well plates filled with 1 mL of LB medium supplemented with the appropriate antibiotic(s) and additives indicated in the text. Bacterial growth, mCherry fluorescence (l_excitation_/l_emission_ = 567 nm/610 nm) and msfGFP fluorescence (l_excitation_/l_emission_ = 475 nm/510 nm) were measured along the experiment in a Synergy HI plate reader (BioTek Instruments, Inc., Winooski, VT, USA). Unless stated otherwise, cultivations were performed in 96-well microtiter plates with a flat bottom and a lid; basal fluorescence levels of *P. putida* KT2440 were subtracted from the individual readings. For physiological characterization of engineered strains, growth kinetics were derived from OD_600_ readings with the light path correction in an ELx808 plate reader (Buch and Holm A/S, Herlev, Denmark) under constant shaking (Wirth *et al*., 2023).

**Table 1.**
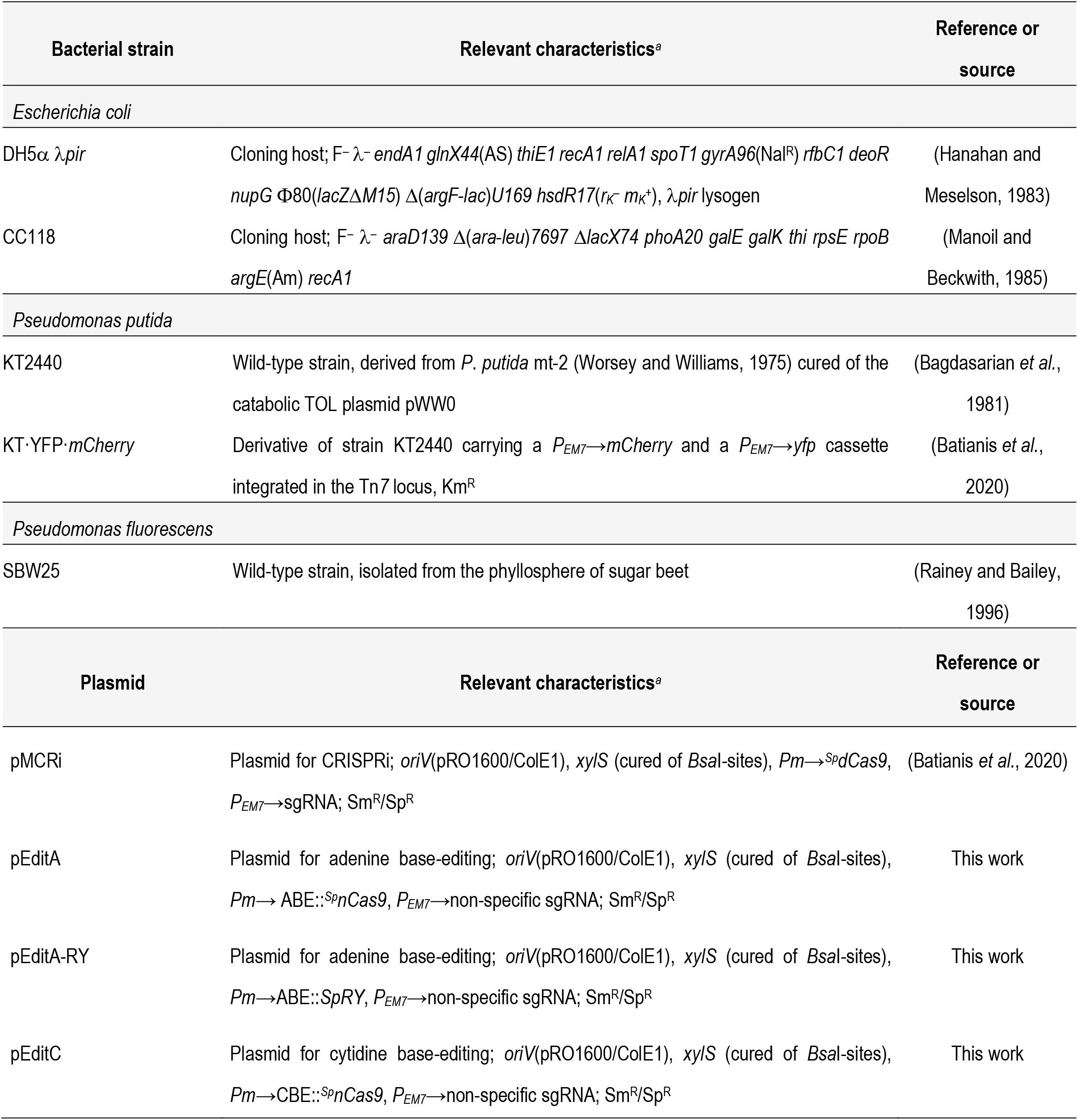
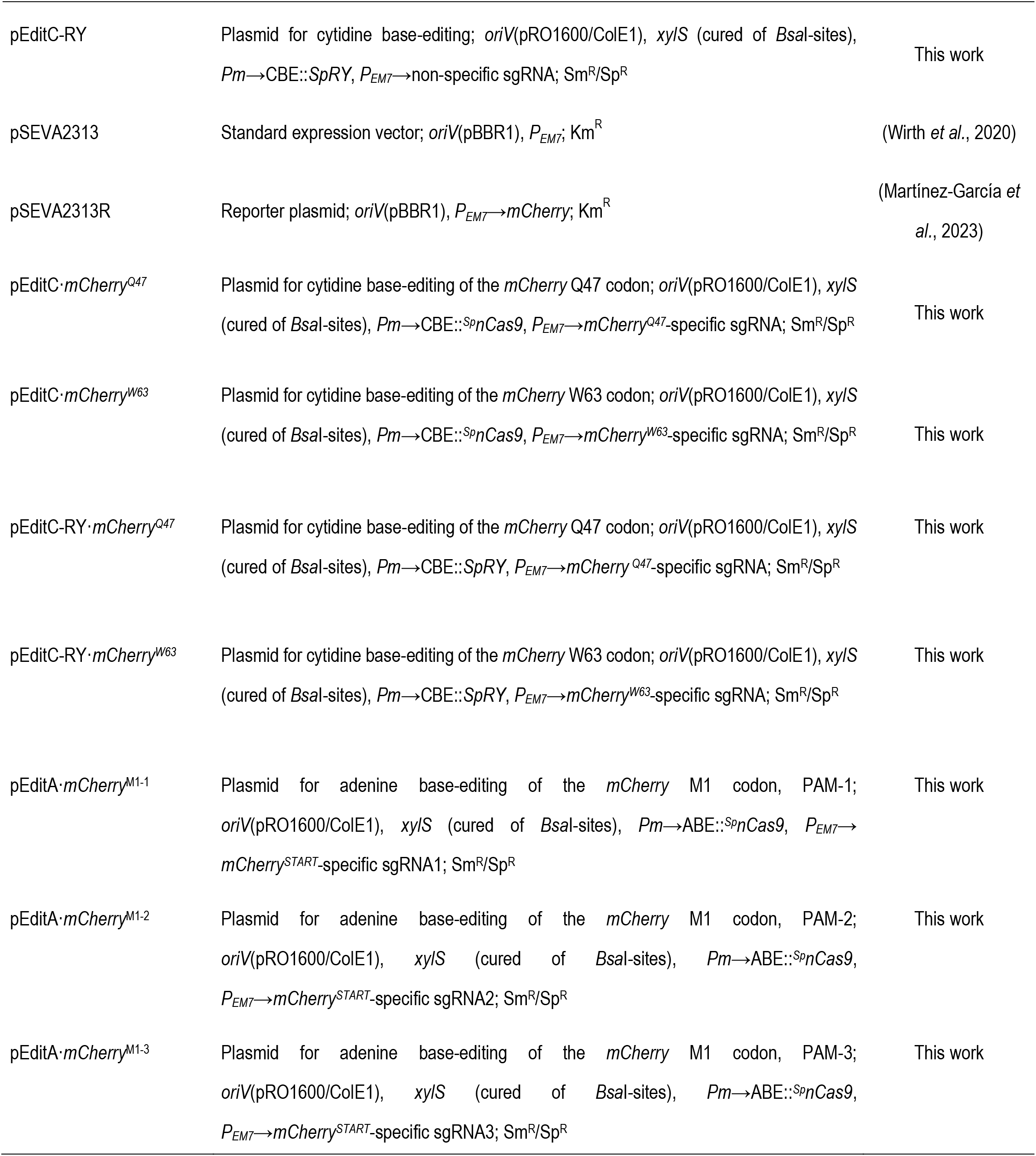
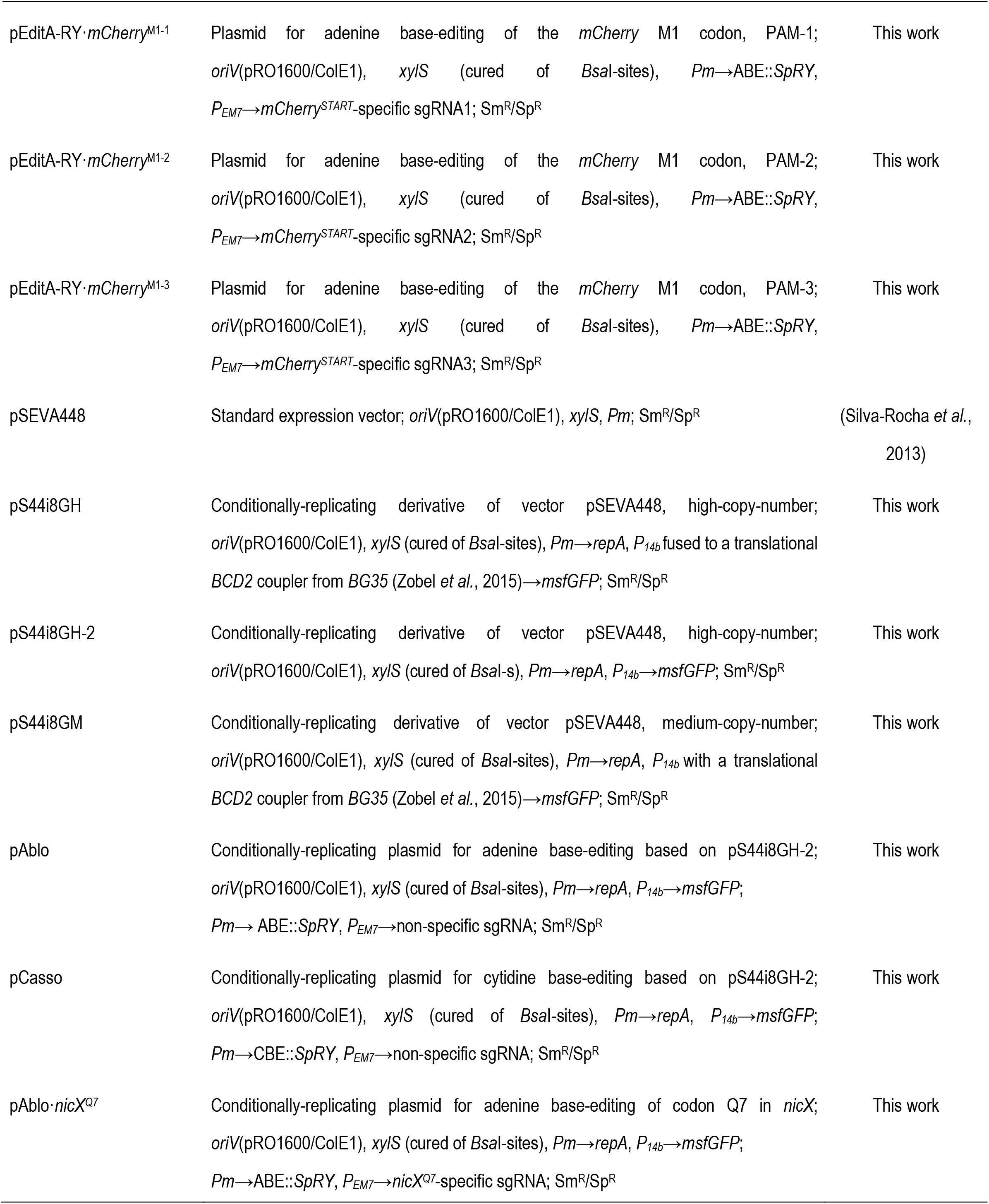
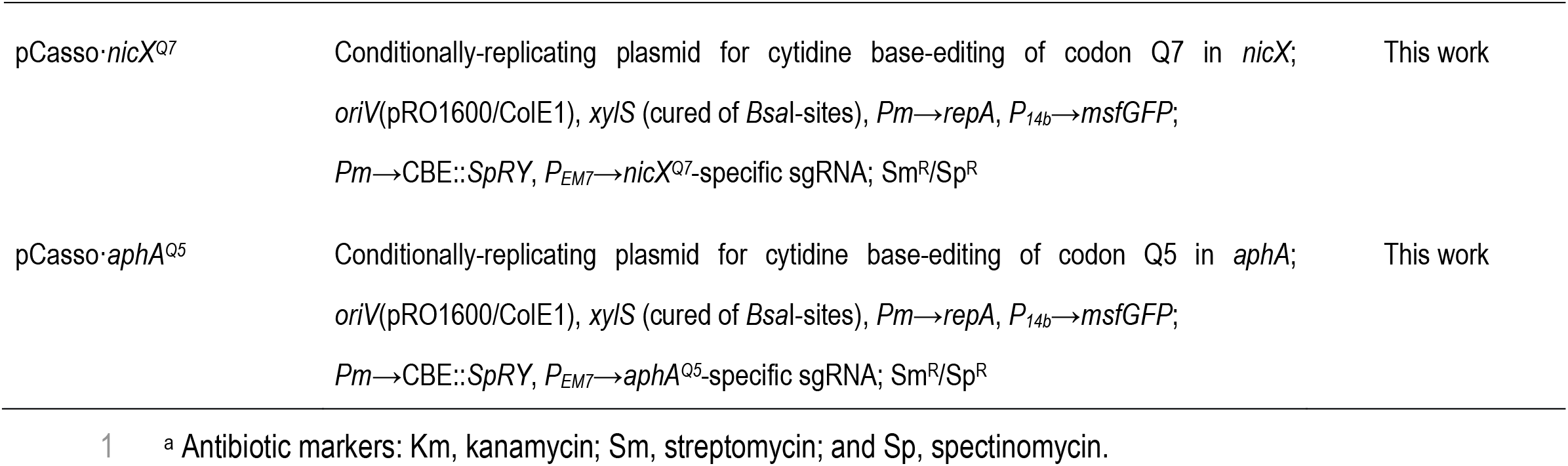
Bacterial strains and plasmids used in this study.

### General cloning procedures and construction of plasmids

Oligonucleotides used in this work are listed in **Supplementary Table S1**. Unless stated otherwise, uracil-excision (*USER*) cloning (Cavaleiro *et al*., 2015) was used for constructing recombinant plasmids; the *AMUSER* tool was employed for designing oligonucleotides (Genee *et al*., 2015). For the construction of the conditionally-replicating plasmids pS44i8GM and pS44iGH, the native promoter and regulatory regions upstream the *repA* gene were inspected *in silico* using BPROM from SoftBerry Inc. (Mount Kisco, NY, USA) (Mohan and Phale, 2017) and PromoTech (Chevez-Guardado and Peña-Castillo, 2021) by following specific instructions for each software package. Phusion™ *U* high-fidelity DNA polymerase (ThermoFisher Scientific) was used according to the manufacturer’s specifications in PCR amplifications intended for *USER* cloning. The *OneTaq*™ master mix (New England BioLabs, Ipswich, MA, USA) was used for most of colony PCRs according to the supplier’s instructions. *E. coli* DH5α λ*pir* (**Table 1**) was employed as a host for cloning purposes. Chemically-competent *E. coli* cells were prepared and transformed with plasmids using the *Mix and Go*™ kit (Zymo Research, Irvin, CA, USA) according to the manufacturer’s indications. Electrocompetent *P. putida* cells were prepared by washing the biomass from saturated LB medium cultures with 300 mM sucrose (Pankratz *et al*., 2023), followed by plasmid transformation through electroporation (Batianis *et al*., 2020). The sequence of all vectors and plasmids used in this study was verified by Mix2Seq sequencing (Eurofins Genomics, Ebersberg, Germany).

### Efficiency of base-editing events through Sanger sequencing of target loci

Bacterial colonies were randomly picked for each condition as described in the corresponding figure legends, and the targeted DNA region was amplified by PCR using the biomass as the template with the Q5™ High-Fidelity 2× master mix (New England Biolabs). Base-editing efficiency when targeting a plasmid-borne gene was estimated by isolating plasmid DNA upon 24-h editing treatment by using the *NucleoSpin*™ plasmid EasyPure kit (Macherey-Nagel, Dueren, Germany). Either the isolated plasmids or the purified PCR products from independent samples were Sanger-sequenced using the *Mix2Seq*™ kit (Eurofins Scientific, Luxembourg) with an adequate pair of oligonucleotides (listed in **Supplementary Table S1**); the resulting DNA sequences were analyzed and visualized using the cloud-based Benchling platform (Benchling Inc., San Francisco, CA, USA). BaseEditR (developed by the Moriarity Laboratory at the University of Minnesota, available at https://github.com/MoriarityLab) was employed to analyze the frequency of base-editing events in plasmid-borne targets.

### Plasmid copy number analysis by quantitative PCR (qPCR) and fluorescence-based counting assays

Plasmid copy number analysis was performed essentially according to the protocol of Fernández-Cabezón et al. (Fernández-Cabezón *et al*., 2022). To this end, a qPCR approach was set in a QuantStudio™ 5 Real-Time PCR system (ThermoFisher Scientific). Expression levels of the *aadA* gene, encoding an aminoglycoside adenylyltransferase that mediates Str resistance, were calculated and converted to the copy number of plasmids carrying the gene. The empty pSEVA448 vector (**Table 1**), was used as a reference for these experiments. Plasmid DNA was purified using *NucleoSpin*™ plasmid EasyPure kit (Macherey-Nagel) according to manufacturer’s instructions, and the plasmid suspensions were diluted to 1-10 ng μL^−1^ to construct calibration curves. *P. putida* cells containing different plasmids as indicated in the text were grown for 10 h in 96-well plates until the cultures reached mid-exponential phase (OD_600_ ~ 0.5). Samples were collected from each well, diluted with phosphate-buffered saline (PBS) to contain 1×10^3^ cells per sample and stored at –20°C until the analysis was carried out. All qPCR amplifications were performed with the following conditions: 50°C for 2 min, 95°C for 10 min, followed by 40 cycles consisting of 95°C for 15 s, 60°C for 1 min and 72°C for 30 s. Each experiment included a non-template control (i.e. a mixture of primers with water, and a mixture of primers with the reaction mix but without any added plasmid). At least three technical replicates were run for each biological sample; results were analyzed in *QuantStudio*™ design and analysis software (ThermoFischer Scientific). A comparative 2^−ΔΔCt^ method (Livak and Schmittgen, 2001) was applied to calculate gene copy numbers by comparing ΔΔCt values of the analyzed *P. putida* clones to the ΔΔCt values of calibration curves.

Fluorescence-based assessment of plasmid loss was done with a 50-μL aliquot of each bacterial culture recovered after the (typically, 24-h long) base-editing procedure. The OD_600_ values of these samples was normalized to 0.5 for each strain and the suspension was plated onto LB medium plates supplemented with 100 μg mL^−1^ Str and 0.1-5 mM of 3-methylbenzoate (3-*m*Bz) as an inducer of the XylS/*Pm* system. All plates were covered with aluminum foil and incubated at 30°C for 24 h to allow for sufficient bacterial growth; after cultivation, the total number of colonies in each plate was counted in an automated UVP ColonyDoc-It imaging station (Analytik Jena US LLC, Upland, CA, USA) with a trisection protocol. Non-fluorescent *P. putida* colonies in each of all three zones were further counted with and without blue light in a transilluminator (ThermoFisher Scientific). The efficiency of plasmid curing (ρ) was calculated as the number of non-fluorescent colonies in each zone divided for the total number of visible colonies in the same area.

### Flow cytometry analysis

Flow cytometry analysis was performed in a Miltenyi MACSQuant™ VYB flow cytometer (Miltenyi Biotec Norden AB, Lund, Sweden), using medium-mixing and fast running mode. In these experiments, 1,000-10,000 events were recorded for each sample. Bacterial cells were gated for singlets in exponential phase, and events within the singlet gate were recorded for each well and each condition analyzed. Median fluorescence values of each triplicate (consisting of a minimum of 5,000 events) were calculated as indicated previously (Nikel *et al*., 2022). The FlowLogic™ software (Inivai Technologies, Mentone, Victoria, Australia) was used for analysis and data interpretation.

### Whole-genome sequencing and analysis of off-target effects

Individual *P. putida* colonies were used to inoculate LB medium cultures (10-mL test tubes), and grown overnight with agitation at 30°C. A 2-mL aliquot of these cultures was used to isolate genomic DNA (gDNA) with the *PureLink*™ gDNA mini kit (ThermoFisher Scientific). Library construction, sequencing, data analysis and subsequent data quality control were performed by Novogene Co. Ltd. (Cambridge, United Kingdom) as previously described (Fernández-Cabezón *et al*., 2021). Briefly, gDNA was randomly sheared into short fragments, and the obtained fragments were end-repaired, A-tailed and further ligated with Illumina adapters. The fragments with adapters were PCR-amplified, size selected and purified.

Sequencing was performed using the Illumina NovaSeq™ 6000 PE150 platform to obtain 150-bp paired end reads. The library was checked with a Qubit 4 fluorometer (ThermoFisher Scientific) and *via* real-time PCR for DNA quantification. A 2100 Bioanalyzer instrument (Agilent Technologies Inc., Santa Clara, CA, USA) was used to estimate size distribution. The resulting libraries were pooled and sequenced on an Illumina, according to the effective library concentration and data amount required in each case. DNA sequencing data was aligned with the reference sequences using the Burrows-Wheeler aligner software (Li and Durbin, 2009). The parameters used were mem -t 4 -k 32 -M, and the mapping rate and coverage were counted according to the alignment results. Duplicates were removed by using the Picard set of command line tools (available at https://github.com/broadinstitute/picard). In this context, single nucleotide polymorphism (SNP) refers to the variation in a single nucleotide that may occur at a specific position in the genome, including transitions and transversions of single nucleotides. To reduce the error rate in SNP detection, the results were filtered according to two criteria: (i) the number of support reads for each SNP should be >4 and (ii) the mapping quality of each detected SNP should be >30. Novogene Co. Ltd. used ANNOVAR (Wang *et al*., 2010) for annotation of the detected SNPs.

### Data and statistical analysis

All figures were created using Adobe Illustrator (Adobe Inc., San Jose, CA, USA); graphics and plots were generated using the Prism 9 GraphPad software (Boston, MA, USA). All the experiments reported in this study were independently repeated at least three times with individual biological replicates (as specified in the corresponding figure legend), and mean values of the corresponding parameter ± standard deviation are presented. Data analysis of qPCR and flow cytometry results was assisted by R programming using RStudio and customized flowCore R scripts (Hahne *et al*., 2009); mean values ± standard deviations were exported to Prism 9 GraphPad for visualization. When relevant, the level of significance of differences when comparing results across experiments was evaluated by ANOVA (Bartlett’s test) with a *P* value < 0.01.

### Protocol for base-editing and facilitated plasmid curing in Gram-negative bacteria

The workflow for a single round of base-editing with pAblo or pCasso plasmids (spanning from plasmid assembly to base-editing procedure followed by plasmid elimination) is schematically illustrated in **Fig. 1**. The specific steps needed for cytidine base-editing (i.e. using the pCasso vector) are detailed below as an example of implementation:

i. **Day 1** · The pCasso vector is digested with FastDigest *Eco*31I (or *Bsa*I) according to the manufacturer’s recommendations. The linearized plasmid is purified after digestion by agarose gel purification of the digestion mixture *via* DNA electrophoresis. The fragment corresponding to the pCasso vector (~12 kb) can be purified in high quantities and stored as a concentrated stock solution at –20°C to be used in multiple applications. To boost ligation efficiency with the linearized vector, oligonucleotides can be phosphorylated using a T4 polynucleotide kinase (PNK, see below) or *via* chemical modification during oligonucleotide synthesis.
ii. The two spacer oligonucleotides (each suspended in Milli-Q H_2_O at 100 μM) are phosphorylated and annealed in a thermocycler. The mixture (10-μL total volume) contains 6 μL of water, 1 μL of each oligonucleotide suspension, 1 μL of T4 ligase buffer and 1 μL of T4 PNK (10 U μL^−1^, ThermoFisher Scientific). The reaction is incubated for 30 min at 37°C, 3 min at 95°C for phosphorylation and T4 PNK deactivation, followed by annealing cycles starting at 95°C and decreasing the temperature by 1°C in each cycle down to 25°C.
iii. The resulting spacer duplex obtained in (ii) is ligated to the linearized pCasso vector. The mixture of double-stranded DNA should be diluted 1:200 with Milli-Q H_2_O. Then, a 10-μL reaction is prepared by mixing 5 μL of the diluted duplex suspension, 1 μL (10 ng) of linearized pCasso vector, 1 μL of T4 ligase buffer, 1 μL of T4 DNA ligase (5 U μL^−1^, ThermoFisher Scientific) and 2 μL of Milli-Q H_2_O (adjusted as needed to reach the total volume). The resulting mixture is incubated for 30 min at room temperature.
iv. A 100-μL aliquot of chemically-competent *E. coli* DH5α cells is transformed with the 10-μL ligation mixture obtained in (iii). The bacterial suspension is plated onto LB medium plates supplemented with Str at 100 μg mL^−1^.
v. **Day 2** · The mini-prep–purified plasmid from 2-3 individual transformants is verified by DNA sequencing with primers 1 and 2 (**Supplementary Table S1**).
vi. **Day 3** · Once the target-specific vectors are constructed and sequence-verified, they are transformed in the recipient host (e.g. *P. putida*) for base-editing. After electroporation of the recombinant pCasso plasmid and recovery of the cells in LB medium at 30°C for 4-16 h, a 10-μL aliquot of the bacterial suspension is inoculated in 10 mL of selective medium (LB medium with 2 mM of 3-*m*Bz and 200 μg mL^−1^ Str) and grown overnight at 30°C. Optionally, the bacterial population could be subjected to passaging into fresh selective medium every 4 h to boost editing efficiency (diluting the culture 1:100 in each passage). Aliquots of the bacterial suspension are plated onto selective medium.
vii. **Day 4** · Editing of the target locus is verified at this stage by picking 10-15 individual colonies (displaying an msfGFP^+^ phenotype), followed by PCR amplification of the target with appropriate oligonucleotides and DNA sequencing.
viii. **Day 5** · Upon confirming the genome modification event(s), the base-editing plasmid is cured from the cells. An individual colony (carrying the intended modification) is streaked onto non-selective medium (LB medium without antibiotic or inducer). After an ~8-16 h incubation at 30°C, the cured (non-fluorescent) colonies can be peaked in a blue-light transilluminator and inoculated for the next round of editing or kept for further strain characterization.

**Figure 1.**
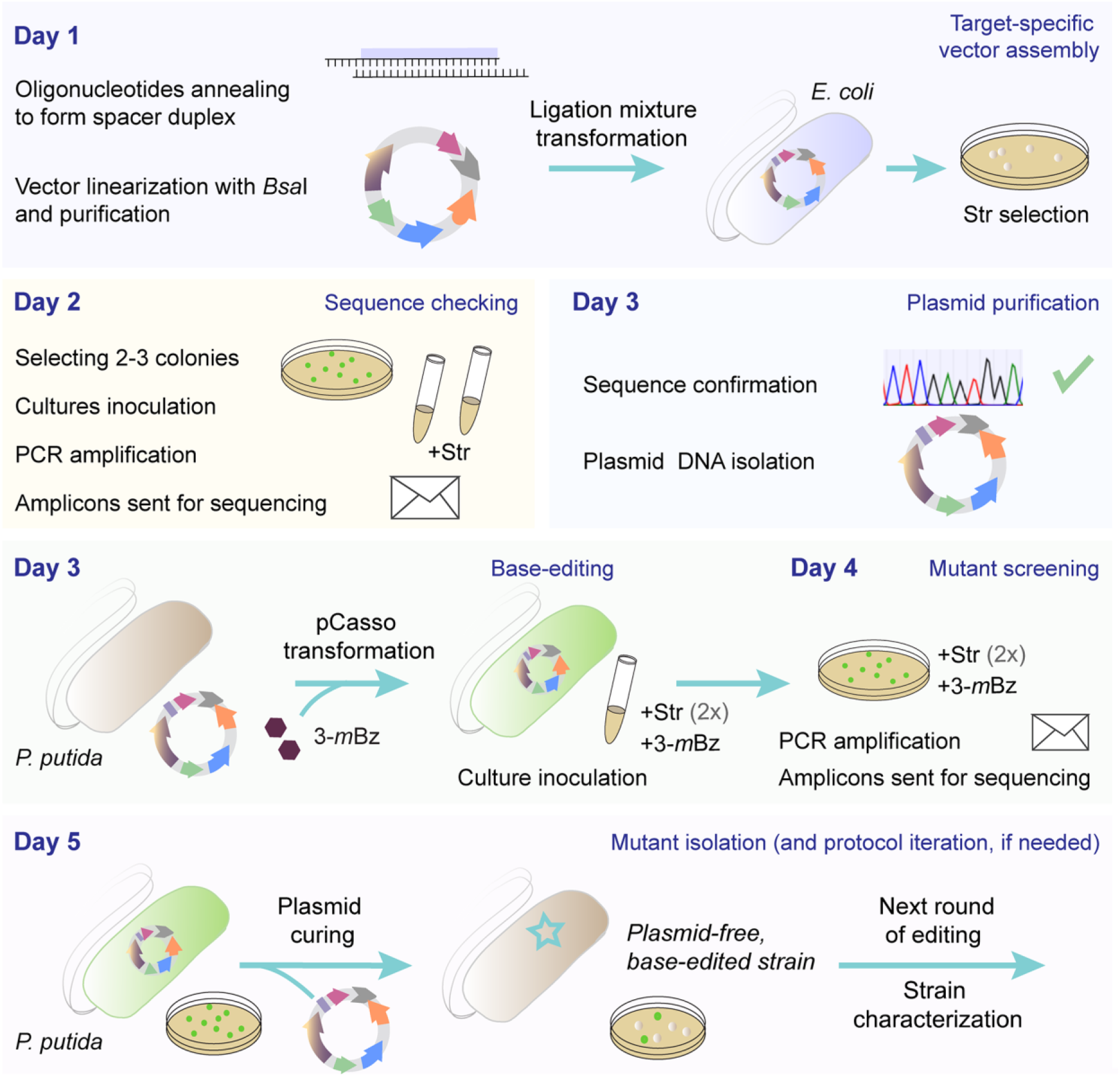
A 5-day workflow for unconstrained cytidine and adenine base-editing in *Pseudomonas*. **Day 1:** single-stranded DNA oligonucleotides are annealed by reverse cooling to form a double-stranded DNA spacer-insert array, which is cloned into the base-editing vector of choice (pAblo or pCasso), previously treated with *Bsa*I. A suitable cloning *E. coli* strain is transformed with the ligation mixture, and the LB medium plates (containing 100 μg mL^−1^ streptomycin for selection, Str) are incubated overnight at 37°C. **Day 2:** 2-3 *E. coli* transformants are inoculated in liquid LB medium with Str, and the cultures are grown overnight at 37°C. PCR amplification and sequencing are used for spacer verification. **Day 3:** sequence-verified plasmids are isolated and transformed into *Pseudomonas* cells by electroporation. Next, a 10-μL aliquot of the cell suspension is inoculated in 10 mL of LB medium under Str selection (2× indicates Str at 200 μg mL^−1^) with 3-methylbenzoate (3-*m*Bz) at 2 mM to ensure plasmid replication and efficient base-editing. Upon recovery at 30°C for 4-16 h, the cell suspension is plated onto LB medium plates added with 2× Str and 2 mM 3-*m*Bz. **Day 4:** green fluorescent colonies (msfGFP^+^) are used as the template for PCR amplification of the target locus. At least 10-15 individual colonies should be checked at this stage, and the corresponding amplicons are sent for sequencing for base-editing verification. **Day 5:** the selected *Pseudomonas* colonies are restreaked by dilution onto non-selective medium (LB medium plates) for plasmid curing (~8-16 h). The cured (non-fluorescent) colonies can be picked under blue light in a transilluminator and inoculated for the next round of base-editing or for strain characterization.

## RESULTS AND DISCUSSION

### Tailoring cytidine and adenine base-editors for precise nucleotide substitutions in bacteria

The use of *^Sp^*Cas9 variants free of PAM restrictions emerged as a powerful option for base-editing in both fundamental and applied research due to significantly improved target accessibility. A version of the *^Sp^*Cas9 nickase (termed SpRY), engineered through structure-guided design, enabled nearly PAM-less genome engineering in human cells (Walton *et al*., 2020), zebrafish (Liang *et al*., 2022), plants (Ren *et al*., 2021) and yeast (Evans and Bernstein, 2021). In this work, SpRY was adopted as the basis to tailor a set of base-editors towards pushing the PAM-restriction boundaries for base-editing in bacteria. Reported to recognize a wide range of PAM sequences, SpRY contains 11 specific amino acid changes as compared to the wild-type *^Sp^*Cas9 nuclease (**Supplementary Table S2**). Recognition of alternative PAMs by SpRY enables targeting DNA sequences with a preference towards 5’-*N*R*N*-3’ over 5’-*N*Y*N*-3’ motifs, where *N* stands for any nucleotide, R is guanine or adenine and Y represents cytosine or thymine (Walton *et al*., 2020). Interestingly, SpRY remains similarly efficient towards the canonical 5’-*N*GG-3’ PAM (**Fig. 2**). This PAM-flexible *^Sp^*Cas9 variant can be useful for locus-restricted base-editing (Anzalone *et al*., 2020). Hence, a CBE, which allows for cytidine (C•G) to thymine (T•A) substitutions (**Fig. 2A**), and an ABE, which converts adenine (A•T) to guanine (G•C) (**Fig. 2B**), were adopted in this study. A CBE typically encompasses three components (Komor *et al*., 2016), i.e. a cytidine deaminase (APOBEC1), fused to a catalytically-impaired *^Sp^*Cas nickase (*^Sp^*nCas9) and two copies of the gene encoding an uracil glycosylase inhibitor (UGI) from *Bacillus subtilis* bacteriophage PBS1, which inhibits counterproductive DNA repair processes (Komor *et al*., 2017; Volke *et al*., 2022). The ABE, on the other hand, spans an evolved tRNA adenine deaminase capable of recognizing ssDNA (TadA from *E. coli*) and *^Sp^*nCas9 (Gaudelli *et al*., 2017). To avoid limitations traditionally associated with the narrow base-editing window of ABEs and their generally low base-editing efficiencies, we chose the ABE8.20-m module, encoding an evolved ABE of the ABEmax8 generation (ABEmax, **Fig. 2B**). This synthetic ABE displays increased activity, improved compatibility with *^Sp^*nCas9 and broadened base-editing window. Owing to these enhanced properties, this base-editor has been deemed suitable for therapeutic applications (Gaudelli *et al*., 2020).

**Figure 2.**
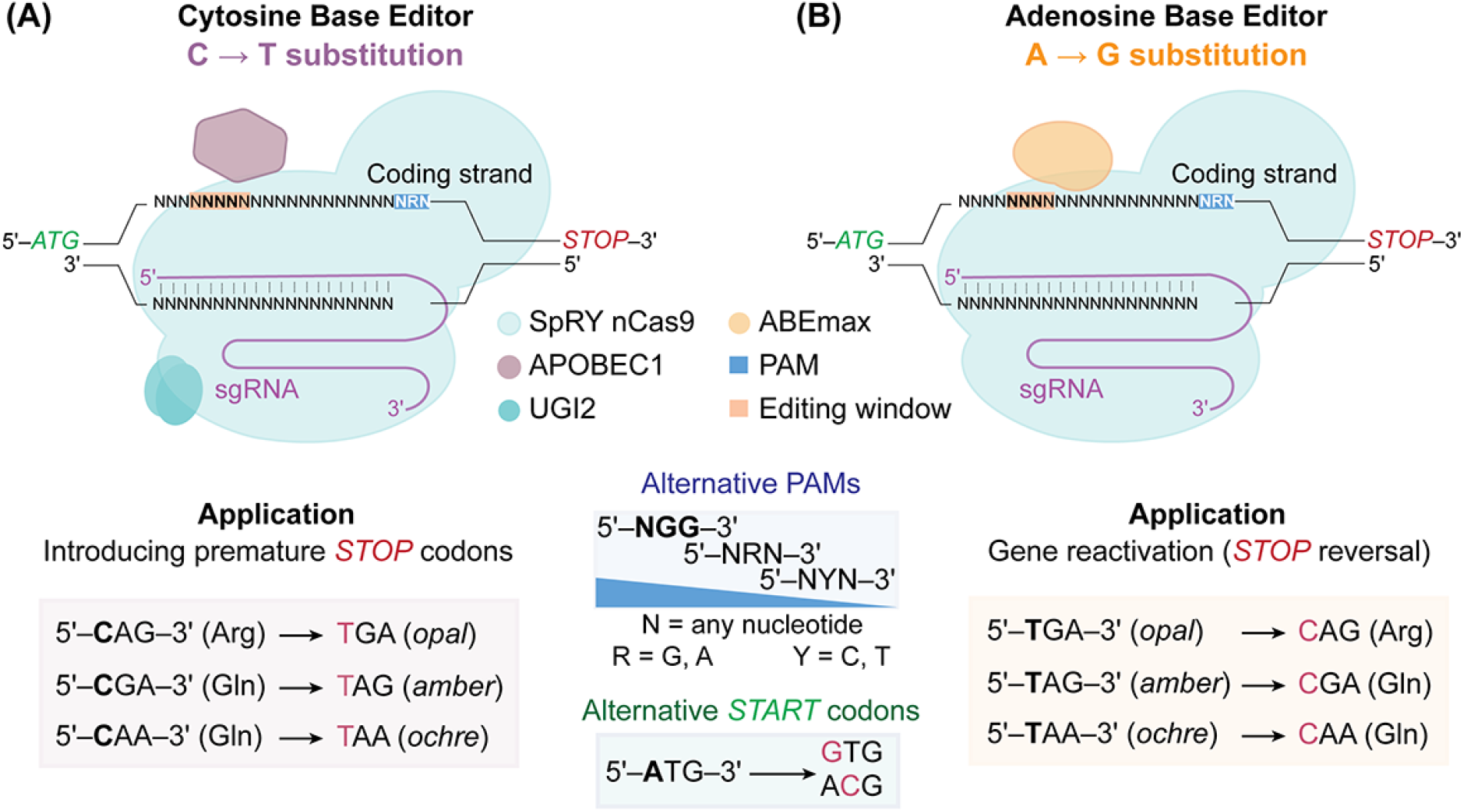
Mechanistic features of the CRISPR/*^Sp^*nCas9 base-editors in this study and their applications for bacterial genome editing. **(A)** Cytidine base-editor. A fusion of SpRY to APOBEC1 (apolipoprotein B mRNA-editing enzyme) and a duplex UGI (uracil DNA glycosylase inhibitor) allows for cytosine-to-thymine substitutions. This editing tool can be exploited for integrating premature *STOP* codon(s) into the reading frame of the gene(s) of interest (generating *opal*, *amber* or *ochre* codons), thus resulting in a functional knock-out. **(B)** Adenine base-editor. In this case, the SpRY variant of the *^Sp^*nCas9 protein is fused to an evolved version of adenine deaminase (ABEmax) that enables adenine-to-guanine substitutions. Application examples of this base-editing tool include restoring gene functionality by modifying premature *STOP* codon(s) back to the wild-type sequence or introducing alternative *START* codons. SpRY allows both base-editors to display a relaxed specificity towards the PAM (protospacer adjacent motif), with a 5’-*N*R*N*-3’ > 5’-*N*Y*N*-3’ preference (where *N* stands for any nucleotide, R is A or G, and Y represents C or T). In both panels, *sgRNA* identifies the synthetic single-guide RNA.

With these functional parts at hand, we built a plasmid-borne set of base-editors as follows. Firstly, all genetic elements of the base-editors previously applied for mammalian cell engineering were repurposed for their use in bacterial systems. DNA fragments encoding SpRY (the nicking variant of *^Sp^*Cas9, carrying an additional D10A mutation), APOBEC1, 2UGI and ABEmax were synthesized *de novo*. The Standard European Vector Architecture (SEVA) is a user-friendly plasmid platform that facilitates standardization in synthetic biology (Silva-Rocha *et al*., 2013). We adopted the SEVA framework, based on swapping and combining individual genetic modules (e.g. promoters, origins of replication and antibiotic markers), to construct the plasmid-born base-editors for bacterial genome engineering. Each of the DNA segments was codon-optimized to facilitate expression in Gram-negative bacteria (Pardo *et al*., 2022) and cured from restriction sites as outlined in SEVA framework. The pMCRi vector series, previously developed in our laboratory for CRISPR interference in *P. putida* (Batianis *et al*., 2020), was used as the physical template of the base-editing toolkit by exchanging the gene encoding *^Sp^*dCas9 (a catalytically inactive Cas9 protein) with the respective base-editor module. All oligonucleotide and gene sequences used to generate these constructs are presented in **Supplementary Tables S1** and **S2**, respectively. These operations gave rise to a first-generation set of SEVA-compatible vectors for base-editing (**Table 1**), i.e. pEditC (carrying a CBE, based on the canonical nicking *^Sp^*nCas9), pEditA (bearing an ABE, based on the canonical nicking *^Sp^*nCas9), pEditC-RY (with a CBE and the evolved nicking SpRY) and pEditA-RY (spanning an ABE and the evolved nicking SpRY). The next step was to test the functionality of these base-editing plasmids in *P. putida*, as indicated in the next section.

### Base-editors enable gene modifications at single-nucleotide resolution both in chromosomally- and plasmid-encoded targets

Cas9-based editing systems require a specific 20-nt spacer sequence in the sgRNA, which determines the navigation of the *^Sp^*nCas9 protein and DNA nicking (i.e. ssDNA break) in the target to be modified. **Fig. 3A** shows an example of a sgRNA sequence binding to the non-coding strand of the DNA encoding the mCherry fluorescent protein. Such a spacer should be selected specifically for each target as a 20-nt sequence followed by a PAM. The canonical PAM for *^Sp^*Cas9 is 5’-*NGG*-3’, and for the engineered SpRY variant, the PAM recognition is relaxed to include the 5’-*N*R*N*-3’ > 5’-*N*Y*N-*3’ variants (**Fig. 2**). An important aspect of the spacer design is to choose unique sequences in the bacterial genome to be edited to reduce off-target effects (**Fig. 3B**). To ensure that this is the case, a simple BLAST analysis (Ladunga, 2017) against the complete gDNA sequence of the bacterial strain at stake should reveal no obvious sequence similarities. In general, the lack of homology in the seed sequence of the sgRNA, comprising 8-10 bases at the 3′-end of the sgRNA-targeting sequence suffices to avoid unintended *^Sp^*nCas9 binding (Wiedenheft *et al*., 2011; Kunne *et al*., 2014). The efficiency of base-editing also depends on the identity of the nucleotides adjacent to the target residue. For instance, the optimal sequence for CBE is 5’-[**T**C ≥ **C**C ≥ **A**C > **G**C]-3’ with the target C nucleotide in the second position, whereas for ABE the preference is 5’-[**T**A ≥ **G**A ≥ **A**A > **C**A]-3’ with the target A nucleotide in the second position (Komor *et al*., 2016; Tong *et al*., 2020).

**Figure 3.**
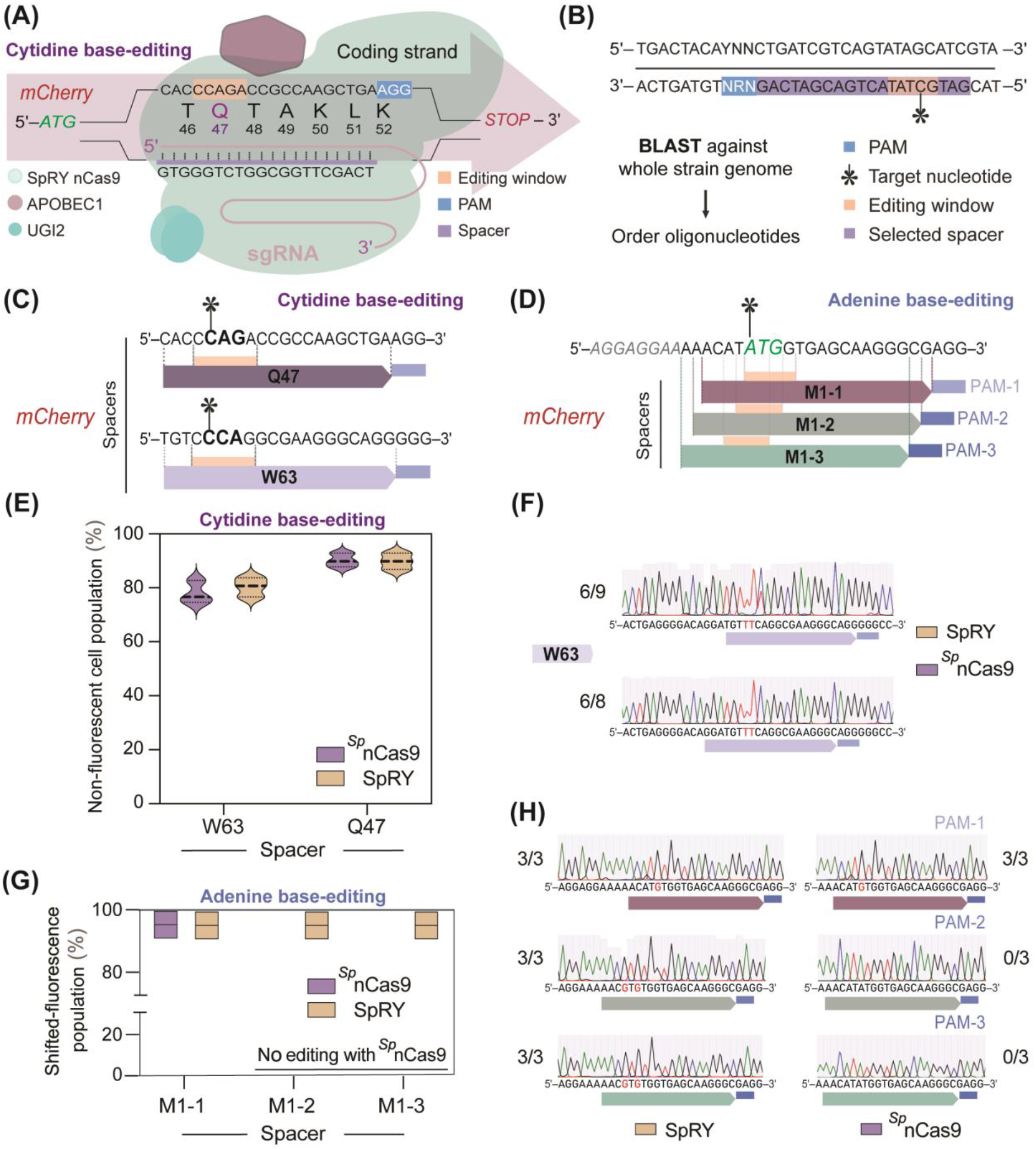
Calibrating cytidine and adenine base-editing in *P. putida*. **(A)** Scheme of an example spacer to edit the cytidine residue within the Q47 codon of the *mCherry* gene; this substitution would result in a premature *STOP* codon. **(B)** Design of target spacers. A PAM sequence (5′-*N*R*N*-3′, where *N* represents any nucleotide and R is either a A or a G) is chosen within the gene of interest, downstream the selected target nucleotide. In this case, the selected spacer is a 20-nt long sequence, which should be validated *via* BLAST against the whole genome to identify potential off-target effects. The optimal editing window is highlighted in orange. **(C)** Cytidine base-editing, targeting codons Q47 and W63 of the *mCherry* gene, with spacers displaying classical (5′-*N*GG-3′) PAMs. **(D)** Adenine base-editing, targeting the *START* codon (M1) of the *mCherry* gene, indicated in green, with spacers displaying classical and non-canonical PAMs. Three spacers were tested to edit the first adenine residue within the *START* codon (highlighted in green). The M1-1 spacer corresponds to 5′-AGG-3′ (i.e. the classical 5′-*N*GG-3′) PAM, while M1-2 and M1-3 are based on 5′-GAG-3′ and 5′-CGA-3′ motifs, respectively (i.e. the alternative, SpRY-compatible 5′-*N*R*N*-3′ PAMs). The Shine-Dalgarno sequence upstream *mCherry* is indicated in gray, italic letters. **(E)** Cytidine base-editing of the *mCherry* gene in *P. putida*. The editing efficiency was estimated after 24 h of treatment using flow cytometry. Strain KT·YFP·*mCherry* (harboring a copy of the *mCherry* gene integrated in the Tn*7* locus of the chromosome) was used as a positive control in these experiments; strain KT2440 was adopted as negative control. Two different spacers were tested to compare the base-editing performance of the canonical *^Sp^*nCas9 and the SpRY variant. **(F)** Base-editing efficiency based on DNA sequencing of the target locus. SpRY had a ~67% editing efficiency (i.e. 6 out of 9 colonies) for two simultaneous editing events (5’-CCC AGA-3’→5’-CT**T** AGA-3’) within the editing window, and *^Sp^*nCas9 displayed a ~75% editing efficiency (i.e. 6 out of 8 colonies) for two or three simultaneous editing events (5’-CCC AGA-3’→5’-C**TT** AGA-3’→5’-**TTT** AGA-3’) within the editing window. **(G)** Adenine base-editing of the *mCherry* gene in *P. putida.* Strains KT·YFP·*mCherry* and KT2440 were used as a positive and negative controls, respectively. *^Sp^*nCas9 with spacers adjacent to non-canonical PAMs mediated no fluorescence shifts (i.e. mCherry levels similar to the positive control, N/A). **(H)** For both *^Sp^*nCas9 and SpRY, DNA sequencing suggested a ~100% editing efficiency (i.e. 3 out of 3 colonies) for 1 editing event (5’-ATG-3’→5’-GTG-3’) using the M1-1 spacer. *^Sp^*nCas9 mediated no detectable base-editing with spacers adjacent to non-canonical PAMs (i.e. M1-2 and M1-3, 0 out of 3 editing colonies). A ~100% editing efficiency was observed for SpRY using M1-2 and M1-3 spacers (3 out of 3 colonies) for two simultaneous editing events (5’-C**A**T **A**TG-3’→5’-C**G**T **G**TG-3’) within the editing window. Experiments indicated in panels **(E)** to **(H)** were independently repeated three times, and randomly-picked colonies were selected for analysis. Nucleotides modified upon base-editing are highlighted in red.

The first-generation vectors in this study (i.e. plasmids pEditA, pEditC, pEditA-SpRY and pEditC-SpRY; **Table 1**) contain two *Bsa*I (*Eco*31I) recognition sites placed upstream of the sgRNA construct. Hence, linearization of the selected plasmid with *Bsa*I allows for the integration of unique target-specific spacers, which can be assembled as a duplex of oligonucleotides complementary to each other (**Fig. 1**). This duplex displays *Bsa*I-compatible overhangs for easy ligation into the destination vector. In particular, one oligonucleotide should contain the 20-nt spacer sequence in the target ORF, flanked with a 5′-GCGCG-3′ motif at the 5′-end; the other oligonucleotide should be its reverse complement with the addition of a 5′-AAAC-3′ sequence to the 5′-end and an extra C nucleotide at the 3’-end. For example, to edit the Q47 codon of *mCherry* with the synthetic CBE (**Fig. 3A** and **3C**), the following oligonucleotides were used to create a duplex *via* direct annealing: CBE_F_*mCherry^Q47^* (5′-GCG CG**C ACC CAG ACC GCC AAG CTG A**-3′) and CBE_R_*mCherry^Q47^* (5′-AAAC **TCA GCT TGG CGG TCT GGG TG**C-3’; primers 3 and 4 in **Supplementary Table S1**). The unique spacer sequence is shown in bold in these oligonucleotides, where the reverse sequence is complementary to the target DNA (**Fig. 3C**). After verifying the plasmid identity and correctness by DNA sequencing, we transformed these vectors into bacteria containing the *mCherry* gene in different configurations to test the base-editors. A similar procedure was adopted to create the spacers used for adenine base-editing (**Fig. 3D**).

To compare the performance of the *^Sp^*nCas9- or SpRY-based CBE in *P. putida,* we used spacers with canonical PAMs targeting the *mCherry* gene (**Fig. 3C**). In particular, the C residues of codons Q47 and W63 were targeted towards creating premature *STOP* codons, thus blocking translation (**Fig. 3A** and **C**). *P. putida* strain KT·YFP·*mCherry* (**Table 1**), previously constructed in our laboratory (Batianis *et al*., 2020), expresses both the *YFP* and *mCherry* genes under control of the synthetic, constitutive *P_EM7_* promoter—conferring yellow and red fluorescence signals that can be used as a readout for calibration of the base-editing toolkit. This reporter system enables simple screening of base-editing events, as the inactivation of the *mCherry* gene through the introduction of a premature *STOP* codon would lead to non-fluorescent bacterial cells. Hence, *P. putida* KT·YFP·*mCherry* was transformed with derivatives of vectors pEditC (i.e. *^Sp^*nCas9-based editor) or pEditC-RY (i.e. SpRY-based editor), endowed with constitutively-expressed sgRNAs carrying target-specific Q47 or W63 spacers adjacent to the canonical 5’-*N*GG-3’ PAM. Specifically, plasmids pEditC·*mCherry^Q47^*, pEditC·*mCherry^W63^*, pEditC-RY·*mCherry^Q47^* and pEditC-RY·*mCherry^W63^* (**Table 1**) were individually delivered into *P. putida* KT·YFP·*mCherry* by electroporation. After incubating the cells for 1 h, a 1-μL aliquot of the culture was inoculated in 10 mL of LB medium and further incubated at 30°C for 24 h. Next, the culture was diluted with PBS buffer to reach a density of 1×10^3^ cells per sample, and analyzed by flow cytometry based on the chromosome-encoded mCherry fluorescence (**Fig. 3E**); the base-editing efficiency was estimated based on the population-wide shift of red fluorescence values between control experiments and cells transformed with base-editing plasmids (**Supplementary Fig. S1A**). Both SpRY and *^Sp^*nCas9 had a similar base-editing performance for canonical PAMs in these experiments (**Fig. 3E**). The observed shift in fluorescence of the base-edited population resulted spacer-dependent and determined by the spacer sequence. When targeting the Q47 codon, 85-95% of the cells had lost the mCherry signal upon base-editing; whereas the non-fluorescent population accounted for 75-85% of the total when the W63 codon was modified (**Fig. 3E** and **Supplementary Fig. S1A**). To further verify base-editing efficiency, individual bacterial colonies were isolated and randomly picked for DNA sequencing of the target region in the chromosome (W63 codon of *mCherry*). Similarly to the estimations based on mCherry fluorescence, the analysis of the SpRY-treated cells revealed a ~67% editing efficiency (with 6 edited targets out of 9 sequenced colonies, **Fig. 3F**), with two simultaneous nucleotide substitutions within the editing window (5’-CCC AGA-3’→5’-C**TT** AGA-3’) and a *STOP* codon incorporated into the ORF at the intended position. In the case of *^Sp^*nCas9, a ~75% editing efficiency was observed (with 6 edited targets out of 8 sequenced colonies, **Fig. 3F**) for two-three simultaneous events within the editing window (5’-CCC AGA-3’→5’-C**TT** AGA-3’→5’-**TTT** AGA-3’).

Since the chromosomally-integrated *mCherry* could be silenced by using *^Sp^*nCas9 and SpRY CBEs, we were interested in testing base-editing of the same gene target in a multicopy format by repeating the experiments with *P. putida* KT2440 carrying plasmid pSEVA2313R (**Table 1**). This reporter plasmid encodes *P_EM7_*→*mCherry* in a vector backbone carrying the low-copy-number *oriV*(RK2) replicon (Jahn *et al*., 2016; Fernández-Cabezón *et al*., 2022). Strain KT2440/pSEVA2313R was transformed with the base-editor plasmids (**Table 1**) and subjected to the flow cytometry analysis explained above. A clear shift in the red fluorescence levels was observed for all the resulting *P. putida* strains (**Supplementary Fig. S1B**). In all cases, a small fraction of mCherry-free *P. putida* cells could still be detected in these cultures (ca. 8-10%), together with an additional (major) sub-population displaying a very low mCherry signal. These results indicate that the *in situ* plasmid editing occurred with a 40-65% efficiency for both *^Sp^*nCas9 and SpRY, thereby yielding bacterial populations with heterogeneous levels of mCherry fluorescence. Based on these observations, the overall base-editing efficiency for multicopy targets dropped by ca. 20% as compared to single-copy gene modifications, while the activity of the *^Sp^*nCas9 and SpRY was comparable in all experiments. Hence, both proteins have similar efficacy in nucleotide substitution while targeting the classical (5’-*N*GG-3’) PAM motif.

### Base-editing with non-canonical PAM sequences

The next step was to test SpRY activity in *P. putida* using spacers with non-canonical PAMs, which would result in an expanded preference to 5’-*NRN*-3’ motifs (where *N* is any nucleotide and R is either A or G). Building on the results in the previous section, we compared both base-editors in strain KT·YFP·*mCherry*, carrying *mCherry* integrated in the *att*Tn*7* site in the chromosome (Batianis *et al*., 2020). We targeted the *START* codon of *mCherry* using 3 different spacers with three different PAMs (**Fig. 3D**), i.e. M1-1 (5’-AGG-3’, canonical sequence), M1-2 (5’-GAG-3’) and M1-3 (5’-CGA-3’). The expected base substitution(s) would result in an alternative *START* codon [5’-GTG-3’ (Ma *et al*., 2002)] in the *mCherry* coding sequence. Such alternative *START* codons are known to affect transcription strength and protein levels (Cao and Slavoff, 2020), and these sequences have been implemented to study transcription activity in bacteria (Belinky *et al*., 2017).

Base-editing was tested in *P. putida* KT·YFP·*mCherry* transformed with plasmids pEditA and pEditA-RY carrying the adenine base-editors and the M1-1, M1-2 and M1-3 spacers (i.e. pEditA·*mCherry*^M1-1,2,3^ and pEditA-RY·*mCherry*^M1-1,2,3^, **Table 1**). As in previous experiments, 1 μL of the bacterial suspension was inoculated in 10 mL of LB medium, and the cells were grown at 30°C for 24 h. Upon plating the cultures on LB agar plates, single colonies were isolated, randomly picked and used for PCR analysis followed by DNA sequencing. Visual inspection of the plates indicated that most of the colonies retained the red fluorescence phenotype albeit at different levels, while some colonies were non-fluorescent (data not shown). Next, base-editing performance was evaluated by flow cytometry, and a new phenotype was observed for cells edited with the M1-1 spacer (encompassing the canonical 5’-NGG-3’ PAM), for both *^Sp^*nCas9 and SpRY. Interestingly, replacing the *ATG* codon of *mCherry* by the (in principle, weaker) *GTG* codon resulted in a heterogeneous population in all base-edited strains, characterized by a fraction of bacteria altogether devoid of mCherry and sub-populations that accumulated different marker amounts— in some cases, higher than those observed in the positive, non-edited control strain. This observation can be accounted by the Shine-Dalgarno dynamics in relation to translation-initiation, which may improve protein expression and counteract adverse mRNA secondary structures (Cao and Slavoff, 2020). Accordingly, >95% of the bacterial population suffered a shift in fluorescence levels for both *^Sp^*nCas9 and SpRY. In contrast, a fluorescence shift in strains harboring SpRY was observed only when the M1-2 and M1-3 (i.e. 5’-*N*RR-3’) spacers were used. The fraction of the cell population with a shifted mCherry level in these cases (~92-95%) proved comparable to the levels observed with the M1-1 spacer (**Fig. 3G** and **Supplementary Fig. S2**). Flow cytometry analysis confirmed the absence of any *^Sp^*nCas9 activity against the M1-2 and M1-3 spacers (**Supplementary Fig. S2**).

To verify the base-editing events and to estimate their efficiency, individual *P. putida* colonies were randomly picked after editing for sequencing analysis of the target locus. A ~100% editing efficiency was calculated for both *^Sp^*nCas9 and SpRY when using the M1-1 spacer for one event (5’-ATG-3’→5’-GTG-3’) in the editing window (**Fig. 3H**). Likewise, strains edited with the non-canonical M1-2 and M1-3 spacers exhibited a ~100% editing efficiency for the SpRY protein for two simultaneous events (5’-C**A**T **A**TG-3’→5’-C**G**T **G**TG-3’) in the editing window (**Fig. 3H**). Expectedly, colonies treated with *^Sp^*nCas9 in combination with spacers adjacent to non-canonical PAMs (i.e. M1-2 and M1-3) shown no detectable base-editing (0 out of 3 colonies), which supports the flow cytometry data (**Fig. 3G**).

The results so far indicate that SpRY can access non-canonical PAMs for base-editing in *P. putida*, whereas *^Sp^*nCas9 is not able to recognize these motifs. Hence, SpRY is a promising addition to the synthetic biology toolbox for base-editing and other CRISPR-Cas9 applications by rendering more targets in the genome accessible to modification. To provide a robust framework for base-editing, we decided to improve the plasmid toolkit in *P. putida* by engineering inducer-dependent curing as explained below.

### Medium- and high-copy-number vectors with conditional replication allow for efficient, one-step plasmid curing

Plasmid-borne tools for genome editing have a drawback that has been rarely addressed in the literature: upon obtaining the desired genotype, the plasmid carrying the functional elements needed for base-editing needs to be cured from the modified strain. Once the plasmid is eliminated, the engineered strain can be used in downstream applications or subjected to further base-editing using different spacers or editors. Plasmid-curing is typically done by repetitive dilution and passaging of bacterial cells in a culture medium with no selection pressure (e.g. antibiotic), and screening of single colonies isolated on non-selective solid media (Buckner *et al*., 2018). Since this process is tedious and time-consuming, an efficient vector-curing approach was recently developed for *Pseudomonas* (Volke *et al*., 2020b). Based on the conditional control of *trfA* expression, the pQURE system mediates inducer-dependent replication of vectors containing the low-copy-number *oriV*(RK2) replicon. While this approach has been successfully implemented for metabolic engineering (Wirth *et al*., 2022), low-copy-number vectors are not ideal for other applications (e.g. base-editing). In order to expand the set of curable vectors available for *Pseudomonas* species and other Gram-negative bacteria, we engineered synthetic replication control in medium- and high-copy-number plasmids based on the *oriV*(pRO1600) replicon by introducing conditional *repA* expression.

Vector pSEVA448 (Silva-Rocha *et al*., 2013) was used as the template in these experiments, as it served as the backbone for constructing various pEdit plasmids (**Table 1**). These plasmids harbor a hybrid of two origins of vegetative replication (i.e. pRO1600 and ColE1). The narrow-host-range *oriV*(ColE1) replicates in *E. coli*, whereas *oriV*(pRO1600), obtained from a *P. aeruginosa* isolate, supports vector replication in *P. putida* and related species (Silva-Rocha *et al*., 2013). A set of plasmids was constructed, collectively termed pS44i8 (where *4i* indicates inducibility of the corresponding origin of replication), bearing an engineered *oriV*(pRO1600). This synthetic module was designed to contain a 3-*m*Bz– inducible XylS/*Pm* expression system to drive conditional expression of *repA*, which encodes the replication protein (**Fig. 4A**). In spite of its widespread use in bacterial genetics, the 5’-UTR region preceding *repA* has been poorly described (e.g. the presence and identity of promoters and Shine-Dalgarno sequences remains virtually unknown). Hence, the predicted native promoter and regulatory regions was explored using *in silico* tools, and two designs were implemented. The predicted *repA* regulatory region was either left intact in front of the *Pm* promoter or completely replaced by the synthetic XylS/*Pm* module (**Supplementary Table S2**). These constructs were designed to facilitate efficient cloning and stable replication in *E. coli* owing to the *oriV*(ColE1) replicon. Conversely, in the absence of 3-*m*Bz, these plasmids would be rapidly and irreversibly lost in *Pseudomonas*. Furthermore, the vectors were endowed with an *msfGFP* gene, encoding the monomeric superfolder green fluorescent protein, to facilitate the screening of plasmid-free colonies. To confer strong, constitutive expression of the reporter gene, a BG35/14b genetic element (Zobel *et al*., 2015), containing *msfGFP* under transcriptional control of the *P_14b_* promoter and a translational coupler (i.e. BCD2 bicistronic design, spanning two Shine-Dalgarno sequences, **Supplementary Table S2**) was adopted. These manipulations gave rise to the conditionally-replicating, medium- and high-copy-number plasmids pS44i8GM pS44i8GH, respectively (**Fig. 4A**). *In vivo* testing was done by transforming plasmids pSEVA448 (control vector), pS44i8GM (native 5’-UTR region preceding *repA* and the *Pm* promoter, **Supplementary Table S2**) and pS44i8GH (synthetic control of *repA* through the XylS/*Pm* system)] into *P. putida* KT2440. The recombinants were inoculated in a microtiter plate in LB medium containing 100 μg mL^−1^ Str and different inducer concentrations (0.5, 1 and 2 mM 3-*m*Bz). After 10 h of incubation at 30°C, cells reached the mid-exponential phase, and samples were collected from each well, diluted with PBS buffer to 1×10^3^ cells per sample. Quantification of gene expression and relative copy number was performed by qPCR; plasmid copy numbers were estimated through the *aadA* (Str^R^) expression rates.

**Figure 4.**
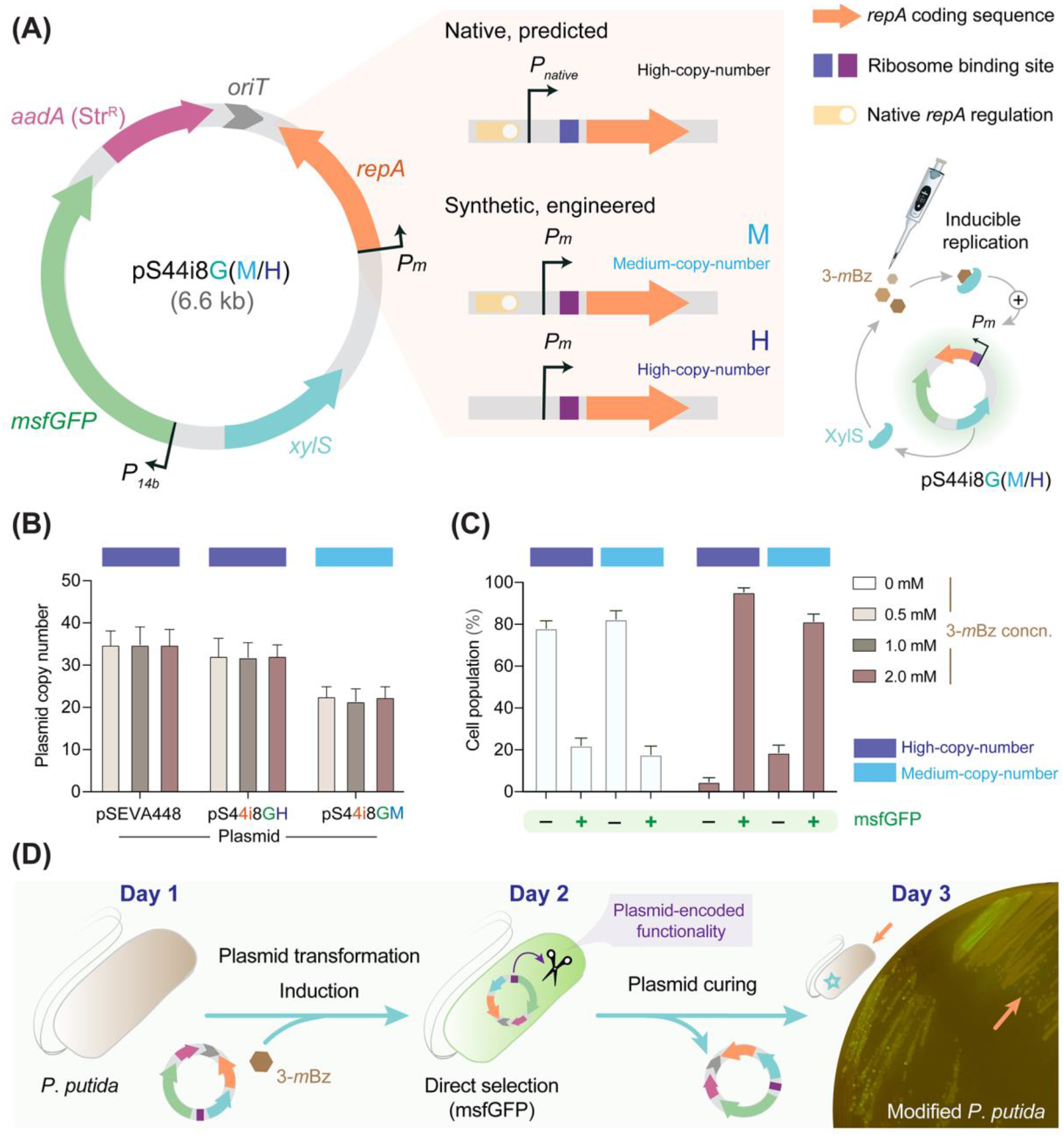
Engineering conditional plasmid replication for *P. putida* based on a high-copy-number vector of the SEVA framework. **(A)** Structure and functional modules of vector pS44i8G; *4i* indicates inducibility of the *oriV*(pRO1600/ColE1) origin of replication, *8* identifies the XylS/*Pm* expression system and *G* stands for *gene of interest* (in the example, *msfGFP*). Integration of a synthetic ribosome binding site and a XylS/*Pm* module replacing the native transcription initiation regulation of *repA* resulted in high-(*H*) and medium-(*M*) copy-number plasmid variants. Further details on relevant genetic elements are indicated in **Table S2**. *Str^R^*, streptomycin-resistance; 3-*m*Bz, 3-methylbenzoate. **(B)** Plasmid stability was analyzed by quantifying the relative copy number of vectors pSEVA448 (control), pS44i8GM (medium-copy-number) and pS44i8GH (high-copy-number) in *P. putida* KT2440 after 10 h of growth (i.e. mid-exponential phase) in LB medium with different inducer concentrations (0.5, 1 and 2 mM 3-*m*Bz). Plasmid copy numbers were calculated by normalizing expression rates measured by qPCR on the plasmid-borne *aadA* gene (Str^R^) and the chromosomally-encoded *rpoB* gene (Fernández-Cabezón *et al*., 2022), and compared with the reference pSEVA448 plasmid (Jahn *et al*., 2016; Martínez-García *et al*., 2023). **(C)** Curing efficiency of vectors pS44i8GM and pS44i8GH as a function of inducer concentration. *P. putida* KT2440 carrying either plasmid was grown for 16 h in LB medium with (0.5, 1 or 2 mM) or without 3-*m*Bz. Cultures were diluted with phosphate-buffered saline to 1×10^3^ cells per sample and analyzed by flow cytometry based on msfGFP fluorescence. The negative (–) population did not display any detectable msfGFP signal (plasmid-free cells); the positive (+) population, in contrast, had different msfGFP levels (plasmid-bearing cells). Experiments in panels **(B)** and **(C)** were independently repeated four times, and mean values ± standard deviations are presented. **(D)** Overview of the plasmid curing approach. Replication of the synthetic *oriV* construct is supported by the presence of 3-*m*Bz; cells transformed with a vector carrying an *8i* module are grown with 1 mM 3-*m*Bz for plasmid maintenance (**Day 1**). On **Day 2**, colonies displaying msfGFP fluorescence are selected, and strain engineering (e.g. base-editing) is carried out in media added with the inducer to sustain vector replication. On **Day 3**, cells are grown in a non-selective medium (i.e. no antibiotic or inducer) and plated onto LB medium *via* dilution-streak to isolate plasmid-free (msfGFP^−^) colonies. Note that conditionally-replicating plasmids can be modified to carry any functionality transiently required for genome editing.

Stable replication was verified both in *E. coli* and *P. putida* for all these plasmids, with copy numbers that remained within the same range regardless of the inducer concentration. No growth on LB medium with Str was registered in control experiments, where no 3-*m*Bz was added to the medium (data not shown). The pS44i8GH vector was maintained at a high-copy number (~32-38 copies per cell), similar to pSEVA448 (~35-42 copies per cell, **Fig. 4B**). These copy number results correlate with data previously reported for SEVA vectors carrying an *oriV*(pRO1600/ColE1) replicon (Jahn *et al*., 2016; Cook *et al*., 2018). Plasmid pS44i8GM, in contrast, consistently exhibited a medium-copy-number (~20-26 copies per cell, **Fig. 4B**). Hence, removing the native regulatory sequences upstream of *repA* resulted in a tighter transcription control of the replication gene. The variations detected in replication efficiency (hence, plasmid copy number) resulted in different levels of msfGFP production, with a ~5-fold decrease in fluorescence levels for *P. putida* harboring plasmid pS44i8GM as compared with pS44i8GH. Building on this concept, the fluorescent marker was exploited for direct screening of cells producing msfGFP by used flow cytometry.

In these experiments, *P. putida* carrying either pS44i8GM or pS44i8GH was grown for 20 h in LB medium with different inducer concentrations (0, 0.5, 1 and 2 mM 3-*m*Bz). The cultures were washed and diluted with PBS buffer to a density of 1×10^3^ cells per sample, followed by flow cytometry analysis based on msfGFP fluorescence. A bimodal distribution was observed in all cases, with a fraction of the cell population displaying an msfGFP^+^ phenotype together with a subpopulation lacking any msfGFP signal, potentially indicating plasmid loss (**Fig. 4C**). When the cells were grown in the presence of 2 mM 3-*m*Bz, >80% of the bacterial population was recovered in the msfGFP^+^ channel for either plasmid tested (**Fig. 4C**). In contrast, when the cells were cultivated in the absence of antibiotic (Str) and inducer of replication (3-*m*Bz), plasmids pS44i8GM (medium-copy-number) and pS44i8GH (high-copy-number) were cured with efficiencies ρ = 84% and 67%, respectively (**Fig. 4C**).

The vectors presented here not only broaden the repertoire of self-curing plasmids available for various manipulations but also offer a choice between medium- and high-copy-number configurations, contingent upon the specific cargo and intended application. These findings underscore the potential of leveraging vector curing as a method for transient activation of plasmid-encoded functions. This approach allows for controlled vector elimination and facilitates screening for the desired phenotypic outcomes in just 3 days, as illustrated in **Fig. 4D**. Guided by this principle, our efforts were directed towards the development of a curable plasmid-based toolkit optimized for near PAM-less base-editing in *P. putida*.

### The pAblo·pCasso toolset for unconstrained base-editing in Gram-negative bacteria

Building on the data above, a pair of self-curing vectors, designated as p**A**blo and p**C**asso (for base-editing of **A**denine and **C**ytidine residues, respectively) were constructed for genome engineering of Gram-negative bacteria. Plasmid pS44i8GH (high-copy-number) was selected as the backbone for integrating the base-editor modules under transcriptional control of the *Pm* promoter. However, preliminary experiments indicated a potentially toxic effect of expressing the base-editor modules together with strong constitutive *msfGFP* expression (data not shown). Hence, the bicistronic design of plasmid pS44i8GH was modified by removing the BCD2 element (Zobel *et al*., 2015), thus keeping a single Shine-Dalgarno sequence (**Supplementary Table S2**). These modifications resulted in plasmid pS448GH-2 (**Table 1**), which was used as the template to add the CBE and ABE modules. The physical map of the resulting pAblo and pCasso vectors (~12 kb) is presented in **Fig. 5A**; the detailed structure of each base-editor module is shown in **Fig. 2**. Considering that the inducer (3-*m*Bz) is needed both for sustaining vector replication (i.e. *Pm*→*repA*) and expression of the base-editor construct (i.e. *Pm*→ABE or CBE), 3-*m*Bz was added to the culture media at 2 mM in all of the experiments described in this section.

**Figure 5.**
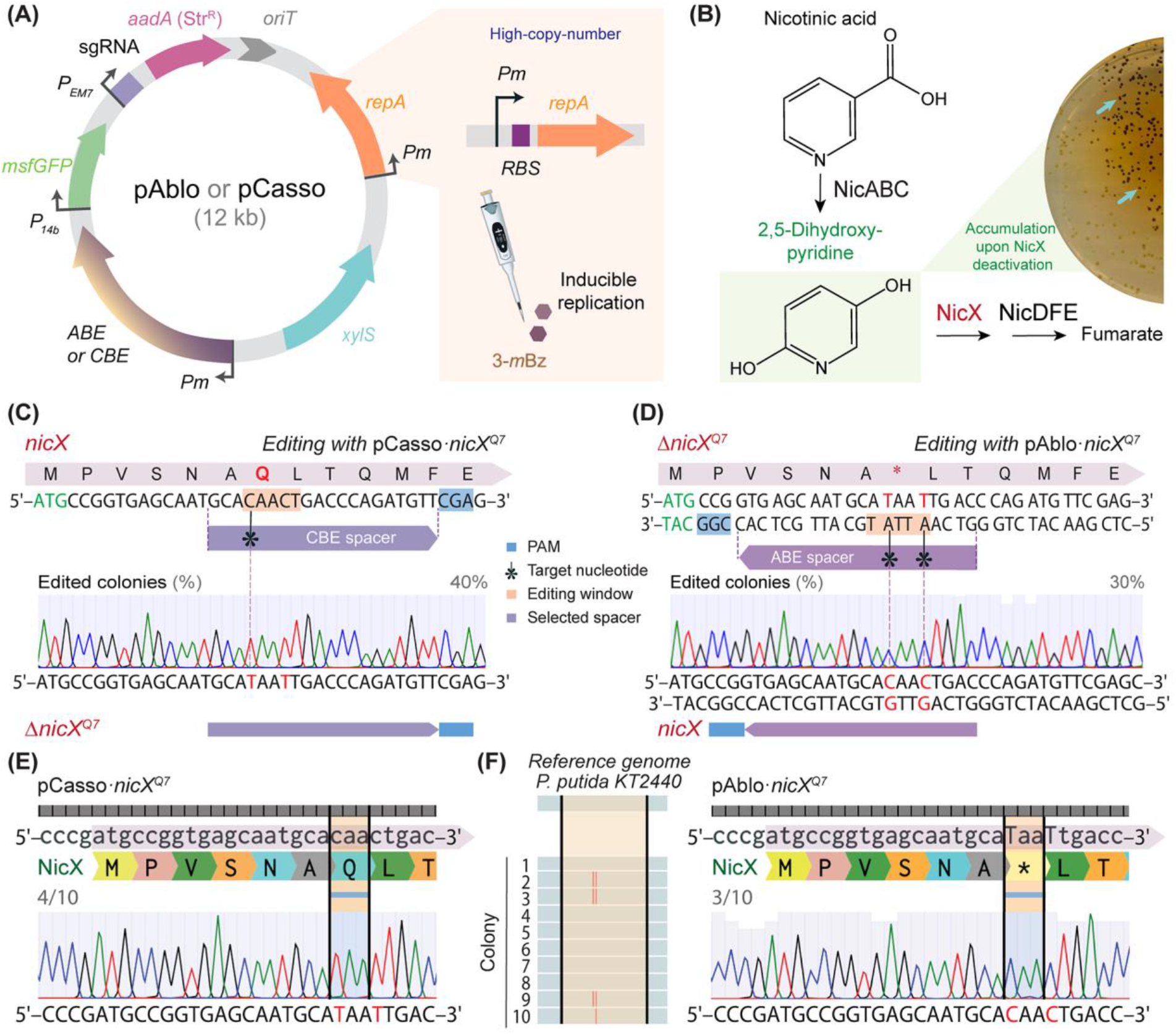
Conditionally-replicating pAblo and pCasso plasmids for base-editing in *P. putida*. **(A)** Scheme of the base-editing pAblo·pCasso plasmids and their functional elements. The high-copy-number pS44i8GH-2 vector (carrying the inducible P*_m_*→*repA* replication module, **Table 1**) was used as backbone; key abbreviations are as follows: *ABE*, adenine base-editor (in pAblo); *CBE*, cytidine base-editor (in pCasso), *msfGFP*, gene encoding the monomeric superfolder green fluorescent protein, *RBS*, ribosome binding site and *sgRNA*, synthetic single-guide RNA. **(B)** Nicotinic acid degradation pathway in *P. putida* KT2440. Functional knock-out of the gene encoding NicX leads to 2,5-dihydroxypyridine accumulation (Wirth *et al*., 2020; Volke *et al*., 2022), which confers a dark-colony phenotype, as illustrated in the photograph (Δ*nicX* colonies incubated for 24 h in LB medium plates containing 5 mM nicotinic acid). **(C)** Base-editing of *nicX* with plasmid pCasso·*nicX*^Q7^ in *P. putida* KT2440. The Q7 glutamine codon (5’-CAA-3’) was targeted to incorporate a premature *STOP* codon (5’-TAA-3’) in the reading frame. PAM, target nucleotide, editing window and selected spacer are highlighted. The percentage of edited colonies was estimated based on the NicX^−^ phenotype indicated in panel **(B)**. **(D)** Base-editing of the Δ*nicX^Q7^* allele with plasmid pAblo·*nicX^Q7^* to restore the wild-type sequence. The *STOP* codon (5’-TAA-3’) was targeted to reverse it back to a glutamine codon (5’-CAA-3’). PAM, target nucleotide, editing window and selected spacer are indicated. **(E)** DNA sequencing indicated a ~40% efficiency (i.e. 4 out of 10 colonies) for two editing events (5’-CAACTG-3’→5’-**T**AA**T**TG-3’), and **(F)** reversion to the original sequence with ~30% efficiency (i.e. 3 out of 10 colonies) for two editing events (5’-TAATTG-3’→5’-**C**AA**C**TG-3’), and 10% (1 out of 10 colonies) for one editing event, not leading to the wild-type sequence restoration (5’-TAATTG-3’→5’-TAA**C**TG-3’). Experiments shown in panels **(C)** to **(F)** were independently repeated three times, and randomly-picked colonies were selected for analysis. Nucleotides modified upon base-editing are highlighted in red.

The functionality of the pAblo·pCasso toolkit was demonstrated by creating a functional gene knockout (i.e. integrating a *STOP* codon into the reading frame, with the pCasso plasmid) followed by restoring its function (i.e. reverting to the original genotype, with the pAblo plasmid). The nicotinic acid degradation pathway of *P. putida* KT2440 (Wirth *et al*., 2020; Volke *et al*., 2022) was selected as a model to evaluate the base-editing capabilities of these tools. Functional knockouts of the *nicX* gene, encoding a 2,5-dihydroxypyridine 5,6-dioxygenase, provide an easy-to-screen phenotype. An interruption of the route at the level of 2,5-dihydroxypyridine leads to the accumulation of this green compound within and outside the colonies, forming a brown polymer upon exposure to oxygen and autoxidation (**Fig. 5B**). Spacers designed for the *nicX* gene were endowed with non-canonical PAMs for targeting the Q7 codon (encoding a glutamine residue, **Fig. 5C** and **Supplementary Table S1**) with both base-editors.

*P. putida* KT2440 was transformed with plasmid pCasso·*nicX^Q7^*, and the resulting strain was grown for 24 h in LB medium with 3-*m*Bz and Str. After plating the bacterial suspension onto LB medium agar containing 5 mM nicotinic acid and incubating the plates for an extra ~16 h, brown colonies accounted for most (>70%) of the bacterial population (**Fig. 5B**). These observations indicate that the CBE borne by pCasso plasmid mediated efficient conversion of cytidine residues in *P. putida*. Next, the base-edited strain was isolated and streaked onto LB agar plates (i.e. without any supplement) to promote plasmid curing. After a ~16 h incubation at 30°C, non-fluorescent colonies were picked by placing the plates onto a blue light transilluminator (i.e. the absence of msfGFP fluorescence under blue light was taken as an indication of plasmid loss, **Fig. 4D**). In the next round of base-editing, we attempted to revert the functional *nicX* knockout in *P. putida* to the original, wild-type genotype. Hence, the base-editing procedure was repeated by transforming vector-free *P. putida* cells (Δ*nicX^Q7^*) with plasmid pAblo·*nicX^Q7^* (**Fig. 5D**). The resulting cultures were incubated for 24 h in LB medium with 3-*m*Bz and Str, and the bacterial suspension was plated onto LB medium agar added with nicotinic acid as explained above. Visual inspection of the plates indicated a wild-type NicX phenotype for roughly half of the *P. putida* colonies. To verify base-editing events in both experiments, 10 individual colonies were randomly picked from the plates for each condition, the target *nicX* locus was amplified by PCR and the amplicons were sequenced. Sequencing analysis confirmed that base-edited colonies represented 40% of the entire pool when using the CBE in plasmid pCasso for introducing a functional *nicX* knockout (**Fig. 5E**). Owing to the editing window of the sequence targeted (**Fig. 5C**), these manipulations led to a double C-to-T modification (i.e. 5’-**C**AA **C**TG-3’→5’-**T**AA **T**TG-3’). Furthermore, sequencing of the amplicons after the second round of base-editing indicated that both nucleotide substitutions were reverted to the original sequence with a ~30% efficiency by using the ABE of plasmid pAblo (**Fig. 5F**).

An important consideration for genome editing is the potential accumulation of off-target mutations (Cho *et al*., 2014). Hence, whole-genome sequencing was adopted to evaluate the frequency of unintended mutations when using CBE and ABE in *P. putida*. Samples were taken after 2 rounds of 24-h base-editing of targets with plasmids pCasso·*nicX^Q7^*, pEditC·*mCherry^W63^*, pAblo·*nicX^Q7^* and pEditA·*mCherry*^M1-1^, and compared to the reference genome (*P. putida* KT2440, grown under the same conditions; **Supplementary Fig. S3**). In general, 12 to 30 off-target mutations were observed in the base-edited isolates. An increased frequency of SNPs was detected in strains treated with SpRY-based editors as compared with *^Sp^*nCas9-based editors (i.e. 17.2% and 18% higher SNP frequency for CBE and ABE, respectively). These values are consistent with off-target effects reported in previous studies using similar base-editors in eukaryotes and other biological systems (Park and Beal, 2019; Cao *et al*., 2022; Wu *et al*., 2022), attributable to the nature of multiple simultaneous base-edits and flexible PAM features of SpRY. No mutations leading to loss-of-function were detected, other than the anticipated introduction of *STOP* codons in target ORFs. Given that the likelihood of spontaneous in-frame mutations that could revert premature *STOP* codons remains minimal even after numerous cell doublings (Volke *et al*., 2022), these observed efficiencies are adequate for microbial strain engineering. This scenario is particularly relevant for generating production strains, proof-of-concept studies and phenotype-based screenings (Wang *et al*., 2023). Based on the results obtained for unconstrained base-editing in *P. putida*, the application of the pAblo·pCasso toolkit was extended to other Gram-negative bacteria as described in the next section.

### Unconstrained base-editing targeting antibiotic resistance determinants in three Gram-negative bacteria

As indicated in previous sections, the pAblo and pCasso plasmids can be used to modify targets encoded either in the bacterial chromosome or in an independent plasmid (**Fig. 3** and **5**, and **Supplementary Fig. S1** and **S2**). To extend the application of nearly-PAMless base-editors, their base-editing performance was analyzed in three Gram-negative bacteria on a multicopy, plasmid-borne target. The *aphA* gene, encoding an aminoglycoside 3′-phosphotransferase, is the antibiotic-resistance determinant in all Km^R^-plasmids of the SEVA framework. *P. putida* KT2440, *P. fluorescens* SBW25 and *E. coli* CC118 were transformed with pSEVA2313R (**Table 1**), which conferred both Km^R^ and mCherry fluorescence in all three bacterial species. These strains were further transformed with plasmid pCasso·*aphA^Q5^*, designed to inactivate *aphA* by introducing a premature *STOP* codon. Cultures were inoculated in LB medium with Str and different inducer concentrations (0, 0.1, 1 and 5 mM 3-*m*Bz for Pseudomonads; 0, 0.1 and 1 mM 3-*m*Bz for *E. coli*; **Fig. 6A**). The OD_600_ of these cultures was followed over 24 h to study the effect of the base-editing system on the overall cell physiology at different levels of induction strength. *P. putida* was not significantly affected by the base-editing platform across all 3-*m*Bz concentrations tested. *P. fluorescens*, in contrast, had longer lag phases and reached lower cell densities under these conditions, with a physiological response dependent on the inducer concentration. At 5 mM 3-*m*Bz, the final OD_600_ values were reduced by ~30%, indicative of a metabolic burden effect (Dvořák *et al*., 2015). The growth of *E. coli* was completely inhibited in the presence of 5 mM 3-*m*Bz, and the final cell density was negatively affected at the inducer concentrations >1 mM (**Fig. 6A**). In spite of the inducer concentration-dependent impact of the pCasso plasmid on growth patterns, we inspected if these treatments mediated the intended *aphA* base-editing event. In this design, the Q5 glutamine codon (5’-CAG-3’) was targeted to introduce a premature *STOP* codon in the *aphA* gene (**Fig. 6B**). *P. putida* colonies were isolated after the base-editing process, and the locus of interest was amplified by PCR and sequenced. All samples tested (4 out of 4 colonies) had the expected base-editing modification in *aphA* (i.e. 5’-CAG-3’→5’-**T**AG-3’), and analysis of the sequencing chromatogram indicated that ~60% of the *aphA* copies have been edited (**Fig. 6B**). The Km-sensitive phenotype was confirmed by streaking the isolated colonies onto LB medium containing the antibiotic (**Fig. 6C**).

**Figure 6.**
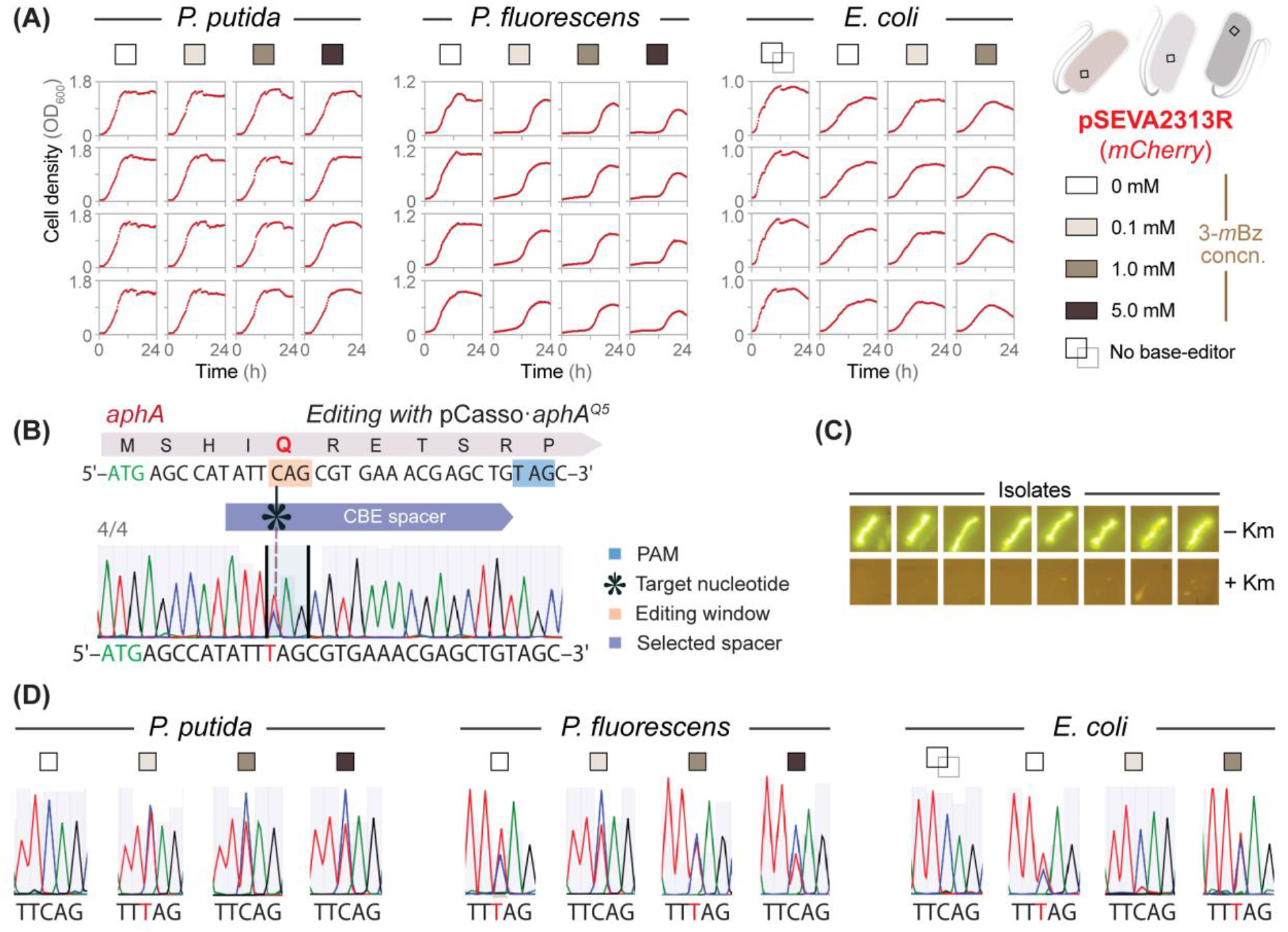
Unconstrained base-editing in Gram-negative bacterial species with conditionally-replicating plasmids. **(A)** Plasmid pCasso·*aphA^Q5^*, targeting *aphA*, an aminoglycoside 3′-phosphotransferase that confers kanamycin (Km) resistance, was used to inactivate the Km-resistance marker borne by plasmid pSEVA2313R in three Gram-negative bacteria (i.e. *P. putida* KT2440, *P. fluorescens* SBW25 and *E. coli* CC118). Cells transformed with both plasmids were grown in LB medium added with different inducer concentrations (0, 0.1, 1 and 5 mM 3-*m*Bz) for 24 h. In control experiments, *E. coli* was transformed with an empty pS448iGH-2 vector (indicated as *No base-editor*). Growth curves are shown in independent quadruplicates for each bacterial species; cell growth was estimated as the optical density measured at 600 nm (*OD_600_*). **(B)** Base-editing of the *aphA* gene with plasmid pCasso·*aphA^Q5^*. The Q5 glutamine codon (5’-CAG-3’) was targeted to incorporate a premature *STOP* codon (5’-TAA-3’) in the reading frame. All samples (4 out of 4 colonies) displayed the correct editing event (5’-CAG-3’→5’-**T**AG-3’), with <50% of all plasmid-borne genes edited as suggested by the DNA sequencing chromatogram. PAM, target nucleotide, editing window and selected spacer are indicated in the diagram. **(C)** Km-sensitivity test for *P. putida* KT2440 carrying the base-edited plasmid pSEVA2313R. Eight randomly-picked colonies were streaked onto LB medium with or without 100 μg mL^−1^ Km and 100 μg mL^−1^ Str (for maintenance of the pCasso·*aphA^Q5^* plasmid). Upon 24 h of incubation at 30°C, plates were photographed under blue light, exploiting the *msfGFP* marker in plasmid pCasso·*aphA^Q5^* as a proxy of bacterial growth. **(D)** DNA sequencing chromatograms to estimate plasmid editing efficiency during pCasso·*aphA^Q5^* base-editing in *P. putida*, *P. fluorescens* and *E. coli*. Independent samples of edited cells were collected in pooled quadruplicates for each condition and analyzed for *aphA* editing events. Nucleotides modified upon base-editing are highlighted in red.

Samples, as pooled quadruplicates for each condition, were also collected to evaluate the plasmid-editing performance by pCasso·*aphA*^Q5^ in all three Gram-negative bacteria. Base-editing events in the *aphA* gene were tested by DNA sequencing of the corresponding amplicons as described above (**Fig. 6D**). While 1 mM 3-*m*Bz was the optimal concentration for plasmid-editing in *P. putida*, a reduced inducer concentration (0.1 mM) demonstrated to be more efficient for multicopy base-editing in *P. fluorescens* and *E. coli* (**Fig. 6D**). A similar editing performance was observed for all three bacterial species, yielding similar results as shown for mCherry modification (**Supplementary Fig. S1B**). Taken together, these results highlight the versatility of the PAM-relaxed variants for base-editing engineering of multiple gene copies in Gram-negative bacteria.

## CONCLUSION

CRISPR-mediated base-editing was introduced in bacteria only recently, yet these technologies have been rapidly adapted for diverse objectives, encompassing both fundamental and applied research (Cho *et al*., 2018; Volke *et al*., 2023). While the PAM requirements of CRISPR systems help mediating sequence-specific adaptive immunity in bacteria, the strict nature of PAM recognition constrains genome engineering applications (Adli, 2018). The SpRY variant circumvents this limitation by relaxing the PAM dependence of *^Sp^*nCas9, which allows for targeting sites with 5’-*N*G*N*-3’ and 5’-*N*A*N*-3’ PAMs—as well as other loci, albeit with reduced recognition efficiency. In this work, an SpRY-based, plasmid-borne toolkit was designed and implemented to break PAM restriction barriers for base-editing of Gram-negative bacterial genomes. We used the nearly PAM-less *^Sp^*nCas9 variant to construct two kinds of base-editors (i.e. ABE and CBE) that enable high-resolution substitutions of virtually any nucleotide in the genome. This is a first-case example of high-efficiency base-editing in Gram-negative bacteria with an evolved version of ABEmax, a tool previously limited to the genome engineering of eukaryotes (Gaudelli *et al*., 2020). Combining ABEmax with the PAM-unrestricted SpRY protein (Walton *et al*., 2020) enabled easy base-editing of multiple DNA sequences. Furthermore, a fast and efficient approach for plasmid curing was implemented, based on conditional vector replication. The resulting plasmids (pS448iGM, pS448iGH and pS448iGH-2) can be equipped with any cargo of choice, which can be subsequently eliminated from the cells in the absence of inducer. Combining this system with the PAM-promiscuous base-editor modules gave rise to the pAblo·pCasso toolset for programmed nucleotide substitution followed by self-curing of the cognate plasmids.

These results demonstrated the functionality of CBE and ABE for genome engineering of *P. putida* and other Gram-negative bacteria—as illustrated by site-directed mutations and their precise reversal in the same engineered strain. The self-curing vectors used for base-editing enable cycling of the genome modifications by means of a simple protocol (**Fig. 1**). Indeed, our results demonstrate that virtually all the cells had lost plasmid pAblo·*nicX*^Q7^ after an 8-h cultivation in LB medium without any additives (ρ = 98%). These functional elements can be adapted for different purposes (including metabolic engineering and exploring gene-to-function relationships) and the base-editing protocol is amenable to high-throughput automation (Gurdo *et al*., 2022; Gurdo *et al*., 2023). The strategy presented in this study eases complex strain engineering programs independently of homologous recombination and yields plasmid-free engineered cells. Furthermore, if crucial for any given application, off-target effects during the base-editing process could be attenuated by employing engineered high-fidelity variants of *^Sp^*Cas9 with improved genome-wide specificities (8). Other modifications could be implemented for easy cloning of target spacers, e.g. in the context of the recently developed Golden Standard (Blázquez *et al*., 2023). In summary, the bacterial genome engineering methodology presented here enables robust constraint-less DNA modifications, paving the way for multiple CRISPR-Cas applications in other bacterial systems (Liu *et al*., 2020).

## Data Availability

The data underlying this article are available in the article and in its supplementary data. All plasmids and data generated in this study are available upon reasonable request. Requests for materials should be addressed to the corresponding author; plasmids pAblo and pCasso can also be obtained from Addgene at https://www.addgene.org.

## Supplementary Data

Supplementary Data are available at NAR Online.

## Authors’ Contributions

**E.K.:** Conceptualization, Investigation, Methodology, Validation, Visualization, Writing – original draft; **Z.S.N.:** Investigation, Data curation, Methodology; **M.N.D.:** Investigation, Data curation, Methodology; **P.I.N:** Conceptualization, Project administration, Supervision, Resources, Funding acquisition, Writing – review & editing.

## Acknowledgements

We are thankful to Daria Sergeeva for technical assistance with qPCR experiments and Mari Rodriguez de Evgrafov (DTU Biosustain) for support with the flow cytometry setup and analysis. Daniel Volke is also gratefully acknowledged for useful comments and handling of materials used in this study. We wish to express our sincere appreciation to the scientific community engaged in metabolic engineering of *Pseudomonas*. Their testing of the toolkit and feedback were instrumental in shaping the outcome of this research.

## Funding

The generous financial support from The Novo Nordisk Foundation through grants NNF20CC0035580, *LiFe* (NNF18OC0034818) and *TARGET* (NNF21OC0067996), and the European Union’s Horizon 2020 Research and Innovation Programme under grant agreement No. 814418 (*SinFonia*) to P.I.N. is gratefully acknowledged. E.K. was supported by the Novo Nordisk Foundation (grant NNF17CC0026768) as part of the Copenhagen Bioscience Ph.D. Programme.

## Conflict of interest statement

None declared.

## Notes

### Competing Interest Statement

The authors have declared no competing interest.

### Summary of Updates

New experimental data added.

