## Supplementary Material for "The pAblo·pCasso self-curing vector toolset for unconstrained cytidine and adenine base-editing in Gram-negative bacteria"

### Supplementary Data

Ekaterina Kozaeva[1], Zacharias S. Nielsen[1], Manuel Nieto-Domínguez[1] and Pablo I. Nikel[1]
[1] The Novo Nordisk Foundation Center for Biosustainability, Technical University of Denmark, 2800 Kgs. Lyngby, Denmark

### Supplementary Tables

**Table S1.** Oligonucleotides used in this work.

| # | Name | Sequence (5'→3') | Application |
|---|---|---|---|
| 1 | Spacer_seq_for | AAAACCCTGGCGACTAGTCTTGGAC | Spacer sequence verification inside of the editing vector |
| 2 | Spacer_seq_rev | AGCAACGTCGGTTCGAGATGGC | |
| 3 | CBE_F_*mCherry*$^{Q47}$ | gcgcgCACCCAGACCGCCAAGCTGA | Spacer cloning for cytidine base-editing of codon Q47 and W63 of *mCherry* |
| 4 | CBE_R_*mCherry*$^{Q47}$ | aaacTCAGCTTGGCGGTCTGGGTGc | |
| 5 | CBE_F_*mCherry*$^{W63}$ | gcgcgTGTCCCAGGCGAAGGGCAGG | |
| 6 | CBE_R_*mCherry*$^{W63}$ | aaacCCTGCCCTTCGCCTGGGACAc | |
| 7 | ABE_F_*mCherry*$^{Q47}$ | gcgcgCGGTCTGGGTGCCCTCGTAG | Spacer cloning for adenine base-editing of codon Q47 and W63 of *mCherry* |
| 8 | ABE_R_*mCherry*$^{Q47}$ | aaacCTACGAGGGCACCCAGACCGc | |
| 9 | ABE_F_*mCherry*$^{W63}$ | gcgcgCCTGGGACATCCTGTCCCCT | |
| 10 | ABE_R_*mCherry*$^{W63}$ | aaacAGGGGACAGGATGTCCCAGGc | |
| 11 | Seq_F_*mCherry*$^{Q47\text{-}W63}$ | CAAGGGCGAGGAGGATAACATGG | *mCherry* sequencing Q47-W63 |
| 12 | Seq_R_*mCherry*$^{Q47\text{-}W63}$ | GCGCAGCTTCACCTTGTAGATGAA | |
| 13 | ABE_F_*mCherry*$^{M1}$-1 | gcgcgACATATGGTGAGCAAGGGCG | Spacer cloning for adenine base-editing of *START* codon M1 of *mCherry* |
| 14 | ABE_R_*mCherry*$^{M1}$-1 | aaacCGCCCTTGCTCACCATATGTc | |
| 15 | ABE_F_*mCherry*$^{M1}$-2 | gcgcgAACATATGGTGAGCAAGGGC | |
| 16 | ABE_R_*mCherry*$^{M1}$-2 | aaacGCCCTTGCTCACCATATGTTc | |
| 17 | ABE_F_mCherry$^{M1}$-3 | gcgcgAAACATATGGTGAGCAAGGG | |
| 18 | ABE_R_*mCherry*$^{M1}$-3 | aaacCCCTTGCTCACCATATGTTTc | |
| 19 | Seq_F_*mCherry*$^{START}$ | CGGCATAGTATATCGGCATAGTA | *mCherry* sequencing M1 |
| 20 | Seq_R_*mCherry*$^{START}$ | AGGCCTTGGAGCCGTACATGAA | |
| 21 | CBE_F_*nicX*$^{Q7}$ | gcgcgGCACAACTGACCCAGATGTT | Spacer cloning for cytidine base-editing of codon Q7 of *nicX* |
| 22 | CBE_R_*nicX*$^{Q7}$ | aaacAACATCTGGGTCAGTTGTGCc | |
| 23 | ABE_F_*nicX*$^{Q7}$ | gcgcgGTCAGTTGTGCATTGCTCAC | Spacer cloning for adenine base-editing of codon Q7 of *nicX* |
| 24 | ABE_R_*nicX*$^{Q7}$ | aaacGTGAGCAATGCACAACTGACc | |
| 25 | CBE_F_*aphA*$^{Q5}$ | gcgcgATTCAGCGTGAAACGAGCTG | Spacer cloning for CBE of codon Q5 of *aphA* |
| 26 | CBE_F_ *aphA*$^{Q5}$ | aaacCAGCTCGTTTCACGCTGAATc | |
| 27 | Seq_F_*aphA* | CACATCGTTCGCCACGCTGCC | *aphA* sequencing (universal Km$^{R}$ gene on SEVA plasmids) |
| 28 | Seq_F_*aphA* | AGTAAACTAGTCTTGGACTCCTGTTG | |
| 29 | Seq_F_*nicX*$^{Q7}$ | AGAGCGTATACGCTACAAATGA | *nicX* sequencing |
| 30 | Seq_R_*nicX*$^{Q7}$ | ATTTCCAGGGTGATGCGCTCGC | |

| | | | |
|---|---|---|---|
| 31 | qPCR_160Str_F | CTTGCAGGTATCTTCGAGC | qPCR fragment amplification for plasmid copy number verification |
| 32 | qPCR_160Str_R | GGCGAGTTCCATAGCGT | |
| 33 | qPCR_80RhoB_F | CAGCAGCCGCTGGGTGGTAA | |
| 34 | qPCR_80RhoB_R | GCGCCGTATGCTTCCAGCG | |

Nucleotides added to the sequences are indicated in blue, lowercase letters.

**Table S2.** Gene fragments and synthetic modules used in this work.

| Description | Sequence (5'→3') |
|---|---|
| Promoter 14b | tTTATTTGACATGCGTGATGTTTAGAATTATAATTTGGGGA |
| Plasmid pS44i8GH [fragment corresponding to promoter 14b followed by the BCD2 bicistronic linker (blue) and *msfGFP* coding sequence]<br><br>*High msfGFP expression* | TTTATTTGACATGCGTGATGTTTAGAATTATAATTTGGGGAcctaggGCCCAAGTTC<br>ACTTAAAAAGGAGATCAACAATGAAAGCAATTTTCGTACTGAAACATCTTAATCATG<br>CTAAGGAGGTtttcta**ATG**ATCATGGGAATTCATAAAGGTGAAGAACTGTTCACCGG<br>TGTTGTTCCGATCCTGGTTGAACTGGATGGTGATGTTAACGGCCACAAATTCTCTGT<br>TCGTGGTGAAGGTGAAGGTGATGCAACCAACGGTAAACTGACCCTGAAATTCATCTG<br>CACTACCGGTAAACTGCCGGTTCCATGGCCGACTCTGGTGACTACCCTGACCTATGG<br>TGTTCAGTGTTTTTCTCGTTACCCGGATCACATGAAGCAGCATGATTTCTTCAAATC<br>TGCAATGCCGGAAGGTTATGTACAGGAGCGCACCATTTCTTTCAAAGACGATGGCAC<br>CTACAAAACCCGTGCAGAGGTTAAATTTGAAGGTGATACTCTGGTGAACCGTATTGA<br>ACTGAAAGGCATTGATTTCAAAGAGGACGGCAACATCCTGGGCCACAAACTGGAATA<br>TAACTTCAACTCCCATAACGTTTACATCACCGCAGACAAACAGAAGAACGGTATCAA<br>AGCTAACTTCAAAATTCGCCATAACGTTGAAGACGGTAGCGTACAGCTGGCGGACCA<br>CTACCAGCAGAACACTCCGATCGGTGATGGTCCGGTTCTGCTGCCGGATAACCACTA<br>CCTGTCCACCCAGTCTAAACTGTCCAAAGACCCGAACGAAAAGCGCGACCACATGGT<br>GCTGCTGGAGTTCGTTACTGCAGCAGGTATCACGCACGGCATGGATGAACTCTACAA<br>A**TAA** |
| Plasmid pS44i8GH-2 [fragment corresponding to promoter 14b followed by a canonical RBS and *msfGFP* coding sequence]<br><br>*Low msfGFP expression* | TTTATTTGACATGCGTGATGTTTAGAATTATAATTTGGGGAccttaaggaggttttc<br>ta**ATG**ATCATGGGAATTCATAAAGGTGAAGAACTGTTCACCGGTGTTGTTCCGATCC<br>TGGTTGAACTGGATGGTGATGTTAACGGCCACAAATTCTCTGTTCGTGGTGAAGGTG<br>AAGGTGATGCAACCAACGGTAAACTGACCCTGAAATTCATCTGCACTACCGGTAAAC<br>TGCCGGTTCCATGGCCGACTCTGGTGACTACCCTGACCTATGGTGTTCAGTGTTTTT<br>CTCGTTACCCGGATCACATGAAGCAGCATGATTTCTTCAAATCTGCAATGCCGGAAG<br>GTTATGTACAGGAGCGCACCATTTCTTTCAAAGACGATGGCACCTACAAAACCCGTG<br>CAGAGGTTAAATTTGAAGGTGATACTCTGGTGAACCGTATTGAACTGAAAGGCATTG<br>ATTTCAAAGAGGACGGCAACATCCTGGGCCACAAACTGGAATATAACTTCAACTCCC<br>ATAACGTTTACATCACCGCAGACAAACAGAAGAACGGTATCAAAGCTAACTTCAAAA<br>TTCGCCATAACGTTGAAGACGGTAGCGTACAGCTGGCGGACCACTACCAGCAGAACA<br>CTCCGATCGGTGATGGTCCGGTTCTGCTGCCGGATAACCACTACCTGTCCACCCAGT<br>CTAAACTGTCCAAAGACCCGAACGAAAAGCGCGACCACATGGTGCTGCTGGAGTTCG<br>TTACTGCAGCAGGTATCACGCACGGCATGGATGAACTCTACAAA**TAA** |
| Native regulatory region of *repA* followed by promoter *Pm* (orange), native RBS and *repA* coding sequence | CAGCTGGGCGCGCCCCTCTCAGGCGCCGCTGGTGCCGCTGGTTGGACGCCAAGGGTG<br>AATCCGCCTCGATACCCTGATTACTCGCTTCCTGCGCCCTCTCAGGCGGCGATAGGG<br>GACTGGTAAAACGGGGATTGCCCAGACGCCTCCCCCGCCCCTTCAGGGGCACAAATG<br>CGGCCCCAACGGGGCCACGTAGTGGTGCGTTTTTTGCGTTTCCACCCTTTTCTTCCT<br>TTTCCCTTTTAAACCTTTTAGGACGTCTACAGGCCACGTAATCCGTGGCCTGTAGAG<br>TTTAAAAAGGGACGGATTTGTTGCCATTAAGGGACGGATTTGTTGTTAAGAAGGGAC<br>GGATTTGTTGTTGTAAAGGGACGGATTTGTTGTATTGTGGGACGCAGATACAGTGTC<br>CCCTTATACACAAGGATAACTAACGTAAggctatctctagtaaggcctaccccttag<br>gctttatgcaacagaaacaataataatggagtcatgaccatgcctagggcgtggacg<br>gcgtcagagaagggagcggacat**ATG**TCGAACGTGGCCTCACCCCCAATGGTTTACA<br>AAAGCAATGCCCTGGTCGAGGCCGCGTATCGCCTCAGTGTTCAGGAACAGCGGATCG<br>TTCTGGCCTGTATTAGCCAGGTGAAGAGGAGCGAGCCTGTCACCGATGAAGTGATGT<br>ATTCAGTGACGGCGGAGGACATAGCGACGATGGCGGGTGTCCCTATCGAATCTTCCT<br>ACAACCAGCTCAAAGAAGCGGCCCTGCGCCTGAAACGGCGGGAAGTCCGGTTAACCC<br>AAGAGCCCAATGGCAAGGGGAAAAGACCGAGTGTGATGATTACCGGCTGGGTGCAAA<br>CAATCATCTACCGGGAGGGTGAGGGCCGTGTAGAACTCAGGTTCACCAAAGACATGC<br>TGCCGTACCTGACGGAACTCACCAAACAGTTCACCAAATACGCCTTGGCTGACGTGG<br>CCAAGATGGACAGCACCCACGCGATCAGGCTTTACGAGCTGCTCATGCAATGGGACA<br>GCATCGGCCAGCGCGAAATAGAAATTGACCAGCTGCGAAAGTGGTTTCAACTGGAAG<br>GCCGGTATCCCTCGATCAAGGACTTCAAGTTGCGAGTGCTTGATCCAGCCGTGACGC<br>AGATCAACGAGCACAGCCCGCTACAGGTGGAGTGGGCGCAGCGAAAGACCGGGCGCA<br>AGGTCACACATCTGTTGTTCAGTTTTGGACCGAAGAAGCCCGCCAAGGCGGTGGGTA<br>AGGCCCCAGCGAAGCGCAAGGCCGGGAAGATTTCAGATGCTGAGATCGCGAAACAGG<br>CTCGCCCTGGTGAGACATGGGAAGCGGCCCGCGCTCGACTAACCCAGATGCCGCTGG<br>ATCTGGCC**TAG** |

| | |
|---|---|
| Modified region of *repA* is followed by promoter *Pm* (orange), native RBS and *repA* coding sequence | CAGCTGGGCGCGCCCCTCTCAGGCGCCGCTGGTGCCGCTGGTTGGACGCCAAGGGTG<br>AATCCGCCTCGATACCCTGATTACTCGCTTCCTGCGCCCTCTCAGGCGGCGATAGGG<br>GACTGGTAAAACGGGGATTGCCCAGACGCCTCCCCCGCCCCTTCAGGGGCACAAATG<br>CGGCCCCAACGGGGCCACGTAGTGGTGCGTTTTTTGCGTTTCCACCCTTTTCTTCCT<br>TTTCCCTTTTAAACCTTTTAGGggctatctctagtaaggcctaccccttaggcttta<br>tgcaacagaaacaataataatggagtcatgaccatgcctagggcgtggacggcgtca<br>gagaagggagcggacat**ATG**TCGAACGTGGCCTCACCCCCAATGGTTTACAAAAGCA<br>ATGCCCTGGTCGAGGCCGCGTATCGCCTCAGTGTTCAGGAACAGCGGATCGTTCTGG<br>CCTGTATTAGCCAGGTGAAGAGGAGCGAGCCTGTCACCGATGAAGTGATGTATTCAG<br>TGACGGCGGAGGACATAGCGACGATGGCGGGTGTCCCTATCGAATCTTCCTACAACC<br>AGCTCAAAGAAGCGGCCCTGCGCCTGAAACGGCGGGAAGTCCGGTTAACCCAAGAGC<br>CCAATGGCAAGGGGAAAAGACCGAGTGTGATGATTACCGGCTGGGTGCAAACAATCA<br>TCTACCGGGAGGGTGAGGGCCGTGTAGAACTCAGGTTCACCAAAGACATGCTGCCGT<br>ACCTGACGGAACTCACCAAACAGTTCACCAAATACGCCTTGGCTGACGTGGCCAAGA<br>TGGACAGCACCCACGCGATCAGGCTTTACGAGCTGCTCATGCAATGGGACAGCATCG<br>GCCAGCGCGAAATAGAAATTGACCAGCTGCGAAAGTGGTTTCAACTGGAAGGCCGGT<br>ATCCCTCGATCAAGGACTTCAAGTTGCGAGTGCTTGATCCAGCCGTGACGCAGATCA<br>ACGAGCACAGCCCGCTACAGGTGGAGTGGGCGCAGCGAAAGACCGGGCGCAAGGTCA<br>CACATCTGTTGTTCAGTTTTGGACCGAAGAAGCCCGCCAAGGCGGTGGGTAAGGCCC<br>CAGCGAAGCGCAAGGCCGGGAAGATTTCAGATGCTGAGATCGCGAAACAGGCTCGCC<br>CTGGTGAGACATGGGAAGCGGCCCGCGCTCGACTAACCCAGATGCCGCTGGATCTGG<br>CC**TAG** |
| Sequence of the *SpRY* gene | **ATG**GACAAGAAGTACAGCATCGGCCTGGATATCGGCACCAACTCTGTGGGCTGGGCC<br>GTGATCACCGACGAGTACAAGGTGCCCAGCAAGAAATTCAAGGTGCTGGGCAACACC<br>GACCGGCACAGCATCAAGAAGAACCTGATCGGAGCCCTGCTGTTCGACAGCGGCGAA<br>ACAGCCGAGAGAACCCGGCTGAAGAGAACCGCCAGAAGAAGATACACCAGACGGAAG<br>AACCGGATCTGCTATCTGCAAGAGATCTTCAGCAACGAGATGGCCAAGGTGGACGAC<br>AGCTTCTTCCACAGACTGGAAGAGTCCTTCCTGGTGGAAGAGGATAAGAAGCACGAG<br>CGGCACCCCATCTTCGGCAACATCGTGGACGAGGTGGCCTACCACGAGAAGTACCCC<br>ACCATCTACCACCTGAGAAAGAAACTGGTGGACAGCACCGACAAGGCCGACCTGCGG<br>CTGATCTATCTGGCCCTGGCCCACATGATCAAGTTCCGGGGCCACTTCCTGATCGAG<br>GGCGACCTGAACCCCGACAACAGCGACGTGGACAAGCTGTTCATCCAGCTGGTGCAG<br>ACCTACAACCAGCTGTTCGAGGAAAACCCCATCAACGCCAGCGGCGTGGACGCCAAG<br>GCCATCCTGTCTGCCAGACTGAGCAAGAGCAGACGGCTGGAAAATCTGATCGCCCAG<br>CTGCCCGGCGAGAAGAAGAATGGCCTGTTCGGAAACCTGATTGCCCTGAGCCTGGGC<br>CTGACCCCCAACTTCAAGAGCAACTTCGACCTGGCCGAGGATGCCAAACTGCAGCTG<br>AGCAAGGACACCTACGACGACGACCTGGACAACCTGCTGGCCCAGATCGGCGACCAG<br>TACGCCGACCTGTTTCTGGCCGCCAAGAACCTGTCCGACGCCATCCTGCTGAGCGAC<br>ATCCTGAGAGTGAACACCGAGATCACCAAGGCCCCCCTGAGCGCCTCTATGATCAAG<br>AGATACGACGAGCACCACCAGGACCTGACCCTGCTGAAAGCTCTCGTGCGGCAGCAG<br>CTGCCTGAGAAGTACAAAGAGATTTTCTTCGACCAGAGCAAGAACGGCTACGCCGGC<br>TACATTGACGGCGGAGCCAGCCAGGAAGAGTTCTACAAGTTCATCAAGCCCATCCTG<br>GAAAAGATGGACGGCACCGAGGAACTGCTCGTGAAGCTGAACAGAGAGGACCTGCTG<br>CGGAAGCAGCGGACCTTCGACAACGGCAGCATCCCCCACCAGATCCACCTGGGAGAG<br>CTGCACGCCATTCTGCGGCGGCAGGAAGATTTTTACCCATTCCTGAAGGACAACCGG<br>GAAAAGATCGAGAAGATCCTGACCTTCCGCATCCCCTACTACGTGGGCCCTCTGGCC<br>AGGGGAAACAGCAGATTCGCCTGGATGACCAGAAAGAGCGAGGAAACCATCACCCCC<br>TGGAACTTCGAGGAAGTGGTGGACAAGGGCGCTTCCGCCCAGAGCTTCATCGAGCGG<br>ATGACCAACTTCGATAAGAACCTGCCCAACGAGAAGGTGCTGCCCAAGCACAGCCTG<br>CTGTACGAGTACTTCACCGTGTATAACGAGCTGACCAAAGTGAAATACGTGACCGAG<br>GGAATGAGAAAGCCCGCCTTCCTGAGCGGCGAGCAGAAAAAGGCCATCGTGGACCTG<br>CTGTTCAAGACCAACCGGAAAGTGACCGTGAAGCAGCTGAAAGAGGACTACTTCAAG<br>AAAATCGAGTGTTTTGATAGTGTTGAAATTTCAGGAGTTGAAGATAGATTTAATGCT<br>TCATTAGGTACCTACCATGATTTGCTAAAAATTATTAAAGATAAAGATTTTTTGGAT<br>AATGAAGAAAATGAAGATATCTTAGAGGATATTGTTTTAACATTGACCTTATTTGAA<br>GATAGGGAGATGATTGAGGAAAGACTTAAAACATATGCTCACCTCTTTGATGATAAG<br>GTGATGAAACAGCTTAAACGTCGCCGTTATACTGGTTGGGGACGTTTGTCTCGAAAA<br>TTGATTAATGGTATTAGGGATAAGCAATCTGGCAAAACAATATTAGATTTTTTGAAA<br>TCAGATGGTTTTGCCAATCGCAATTTTATGCAGCTGATCCATGATGATAGTTTGACA<br>TTTAAAGAAGACATTCAAAAAGCACAAGTGTCTGGACAAGGCGATAGTTTACATGAA<br>CATATTGCAAATTTAGCTGGTAGCCCTGCTATTAAAAAAGGTATTTTACAGACTGTA |

| | |
|---|---|
| | AAAGTTGTTGATGAATTGGTCAAAGTAATGGGGCGGCATAAGCCAGAAAATATCGTT<br>ATTGAAATGGCACGTGAAAATCAGACAACTCAAAAGGGCCAGAAAAATTCGCGAGAG<br>CGTATGAAACGAATCGAAGAAGGTATCAAAGAATTAGGAAGTCAGATTCTTAAAGAG<br>CATCCTGTTGAAAATACTCAATTGCAAAATGAAAAGCTCTATCTCTATTATCTCCAA<br>AATGGAAGAGACATGTATGTGGACCAAGAATTAGATATTAATCGTTTAAGTGATTAT<br>GATGTCGATCACATTGTTCCACAAAGTTTCCTTAAAGACGATTCAATAGACAATAAG<br>GTCTTAACGCGTTCTGATAAAAATCGTGGTAAATCGGATAACGTTCCAAGTGAAGAA<br>GTAGTCAAAAAGATGAAAAACTATTGGAGACAACTTCTAAACGCCAAGTTAATCACT<br>CAACGTAAGTTTGATAATTTAACGAAAGCTGAACGTGGAGGTTTGAGTGAACTTGAT<br>AAAGCTGGTTTTATCAAACGCCAATTGGTTGAAACTCGCCAAATCACTAAGCATGTG<br>GCACAAATTTTGGATAGTCGCATGAATACTAAATACGATGAAAATGATAAACTTATT<br>CGAGAGGTTAAAGTGATTACCTTAAAATCTAAATTAGTTTCTGACTTCCGAAAAGAT<br>TTCCAATTCTATAAAGTACGTGAGATTAACAATTACCATCATGCCCATGATGCGTAT<br>CTAAATGCCGTCGTTGGAACTGCTTTGATTAAGAAATATCCAAAACTTGAATCGGAG<br>TTTGTCTATGGTGATTATAAAGTTTATGATGTTCGTAAAATGATTGCTAAGTCTGAG<br>CAAGAAATAGGCAAAGCAACCGCAAAATATTTCTTTTACTCTAATATCATGAACTTC<br>TTCAAAACAGAAATTACACTTGCAAATGGAGAGATTCGCAAACGCCCTCTAATCGAA<br>ACTAATGGGGAAACTGGAGAAATTGTCTGGGATAAAGGGCGAGATTTTGCCACAGT<br>GCGCAAAGTATTGTCCATGCCCCAAGTCAATATTGTCAAGAAAACAGAAGTACAGAC<br>AGGCGGATTCTCCAAGGAGTCAATCCGGCCCAAGCGGAACAGCGATAAGCTCATCGC<br>GCGGAAAAAAGATTGGGACCCTAAAAAGTATGGGGGGTTCTTGTGGCCGACGGTCGC<br>TTACAGCGTCTTGGTCGTGGCAAAGGTCGAAAAGGGCAAGTCGAAAAAGCTCAAATC<br>GGTCAAAGAGTTGTTGGGTATTACGATTATGGAGCGGTCCTCGTTCGAAAAGAACCC<br>CATCGACTTTCTGGAAGCCAAAGGCTATAAGGAAGTCAAGAAGGATCTCATCATCAA<br>ATTGCCGAAATATTCCCTCTTTGAATTGGAAAACGGTCGTAAGCGTATGCTGGCCTC<br>GGCGAAGCAACTGCAGAAAGGGAACGAATTGGCGCTCCCTTCCAAGTATGTGAACTT<br>CCTCTACCTCGCGTCGCACTACGAGAAGTTGAAAGGGTCGCCAGAAGATAATGAACA<br>AAAGCAGCTCTTTGTGGAGCAGCACAAGCATTACCTCGACGAGATTATCGAGCAAAT<br>TAGCGAGTTCAGCAAACGGGTCATTCTCGCCGACGCCAATCTCGACAAAGTGCTGTC<br>CGCATACAATAAGCATCGGGACAAACCAATTCGGGAGCAAGCCGAAAATATCATCCA<br>CCTGTTTACCTTGACCCGCCTGGGTGCCCCTCGCGCGTTCAAGTACTTTGATACCAC<br>CATCGACCCAAAACAGTACCGGTCCACGAAGGAGGTCCTCGATGCTACCCTCATCCA<br>TCAGAGCATTACCGGCCTCTATGAGACGCGCATCGACCTCTCG**TGA** |
| Sequence of the nicking *SpRY* gene | **ATG**GACAAGAAGTACAGCATCGGCCTGGCCATCGGCACCAACTCTGTGGGCTGGGCC<br>GTGATCACCGACGAGTACAAGGTGCCCAGCAAGAAATTCAAGGTGCTGGGCAACACC<br>GACCGGCACAGCATCAAGAAGAACCTGATCGGAGCCCTGCTGTTCGACAGCGGCGAA<br>ACAGCCGAGAGAACCCGGCTGAAGAGAACCGCCAGAAGAAGATACACCAGACGGAAG<br>AACCGGATCTGCTATCTGCAAGAGATCTTCAGCAACGAGATGGCCAAGGTGGACGAC<br>AGCTTCTTCCACAGACTGGAAGAGTCCTTCCTGGTGGAAGAGGATAAGAAGCACGAG<br>CGGCACCCCATCTTCGGCAACATCGTGGACGAGGTGGCCTACCACGAGAAGTACCCC<br>ACCATCTACCACCTGAGAAAGAAACTGGTGGACAGCACCGACAAGGCCGACCTGCGG<br>CTGATCTATCTGGCCCTGGCCCACATGATCAAGTTCCGGGGCCACTTCCTGATCGAG<br>GGCGACCTGAACCCCGACAACAGCGACGTGGACAAGCTGTTCATCCAGCTGGTGCAG<br>ACCTACAACCAGCTGTTCGAGGAAAACCCCATCAACGCCAGCGGCGTGGACGCCAAG<br>GCCATCCTGTCTGCCAGACTGAGCAAGAGCAGACGGCTGGAAAATCTGATCGCCCAG<br>CTGCCCGGCGAGAAGAAGAATGGCCTGTTCGGAAACCTGATTGCCCTGAGCCTGGGC<br>CTGACCCCCAACTTCAAGAGCAACTTCGACCTGGCCGAGGATGCCAAACTGCAGCTG<br>AGCAAGGACACCTACGACGACGACCTGGACAACCTGCTGGCCCAGATCGGCGACCAG<br>TACGCCGACCTGTTTCTGGCCGCCAAGAACCTGTCCGACGCCATCCTGCTGAGCGAC<br>ATCCTGAGAGTGAACACCGAGATCACCAAGGCCCCCCTGAGCGCCTCTATGATCAAG<br>AGATACGACGAGCACCACCAGGACCTGACCCTGCTGAAAGCTCTCGTGCGGCAGCAG<br>CTGCCTGAGAAGTACAAAGAGATTTTCTTCGACCAGAGCAAGAACGGCTACGCCGGC<br>TACATTGACGGCGGAGCCAGCCAGGAAGAGTTCTACAAGTTCATCAAGCCCATCCTG<br>GAAAAGATGGACGGCACCGAGGAACTGCTCGTGAAGCTGAACAGAGAGGACCTGCTG<br>CGGAAGCAGCGGACCTTCGACAACGGCAGCATCCCCCACCAGATCCACCTGGGAGAG<br>CTGCACGCCATTCTGCGGCGGCAGGAAGATTTTTACCCATTCCTGAAGGACAACCGG<br>GAAAAGATCGAGAAGATCCTGACCTTCCGCATCCCCTACTACGTGGGCCCTCTGGCC<br>AGGGGAAACAGCAGATTCGCCTGGATGACCAGAAAGAGCGAGGAAACCATCACCCCC<br>TGGAACTTCGAGGAAGTGGTGGACAAGGGCGCTTCCGCCCAGAGCTTCATCGAGCGG<br>ATGACCAACTTCGATAAGAACCTGCCCAACGAGAAGGTGCTGCCCAAGCACAGCCTG<br>CTGTACGAGTACTTCACCGTGTATAACGAGCTGACCAAAGTGAAATACGTGACCGAG |

| | |
|---|---|
| | GGAATGAGAAAGCCCGCCTTCCTGAGCGGCGAGCAGAAAAAGGCCATCGTGGACCTG<br>CTGTTCAAGACCAACCGGAAAGTGACCGTGAAGCAGCTGAAAGAGGACTACTTCAAG<br>AAAATCGAGTGTTTTGATAGTGTTGAAATTTCAGGAGTTGAAGATAGATTTAATGCT<br>TCATTAGGTACCTACCATGATTTGCTAAAAATTATTAAAGATAAAGATTTTTTGGAT<br>AATGAAGAAAATGAAGATATCTTAGAGGATATTGTTTTAACATTGACCTTATTTGAA<br>GATAGGGAGATGATTGAGGAAAGACTTAAAACATATGCTCACCTCTTTGATGATAAG<br>GTGATGAAACAGCTTAAACGTCGCCGTTATACTGGTTGGGGACGTTTGTCTCGAAAA<br>TTGATTAATGGTATTAGGGATAAGCAATCTGGCAAAACAATATTAGATTTTTTGAAA<br>TCAGATGGTTTTGCCAATCGCAATTTTATGCAGCTGATCCATGATGATAGTTTGACA<br>TTTAAAGAAGACATTCAAAAAGCACAAGTGTCTGGACAAGGCGATAGTTTACATGAA<br>CATATTGCAAATTTAGCTGGTAGCCCTGCTATTAAAAAAGGTATTTTACAGACTGTA<br>AAAGTTGTTGATGAATTGGTCAAAGTAATGGGGCGGCATAAGCCAGAAAATATCGTT<br>ATTGAAATGGCACGTGAAAATCAGACAACTCAAAAGGGCCAGAAAAATTCGCGAGAG<br>CGTATGAAACGAATCGAAGAAGGTATCAAAGAATTAGGAAGTCAGATTCTTAAAGAG<br>CATCCTGTTGAAAATACTCAATTGCAAAATGAAAAGCTCTATCTCTATTATCTCCAA<br>AATGGAAGAGACATGTATGTGGACCAAGAATTAGATATTAATCGTTTAAGTGATTAT<br>GATGTCGATCACATTGTTCCACAAAGTTTCCTTAAAGACGATTCAATAGACAATAAG<br>GTCTTAACGCGTTCTGATAAAAATCGTGGTAAATCGGATAACGTTCCAAGTGAAGAA<br>GTAGTCAAAAAGATGAAAAACTATTGGAGACAACTTCTAAACGCCAAGTTAATCACT<br>CAACGTAAGTTTGATAATTTAACGAAAGCTGAACGTGGAGGTTTGAGTGAACTTGAT<br>AAAGCTGGTTTTATCAAACGCCAATTGGTTGAAACTCGCCAAATCACTAAGCATGTG<br>GCACAAATTTTGGATAGTCGCATGAATACTAAATACGATGAAAATGATAAACTTATT<br>CGAGAGGTTAAAGTGATTACCTTAAAATCTAAATTAGTTTCTGACTTCCGAAAAGAT<br>TTCCAATTCTATAAAGTACGTGAGATTAACAATTACCATCATGCCCATGATGCGTAT<br>CTAAATGCCGTCGTTGGAACTGCTTTGATTAAGAAATATCCAAAACTTGAATCGGAG<br>TTTGTCTATGGTGATTATAAAGTTTATGATGTTCGTAAAATGATTGCTAAGTCTGAG<br>CAAGAAATAGGCAAAGCAACCGCAAAATATTTCTTTTACTCTAATATCATGAACTTC<br>TTCAAAACAGAAATTACACTTGCAAATGGAGAGATTCGCAAACGCCCTCTAATCGAA<br>ACTAATGGGGAAACTGGAGAAATTGTCTGGGATAAAGGGCGAGATTTTGCCACAGTG<br>CGCAAAGTATTGTCCATGCCCCAAGTCAATATTGTCAAGAAAACAGAAGTACAGACA<br>GGCGGATTCTCCAAGGAGTCAATCCGGCCCAAGCGGAACAGCGATAAGCTCATCGCG<br>CGGAAAAAAGATTGGGACCCTAAAAAGTATGGGGGGTTCTTGTGGCCGACGGTCGCT<br>TACAGCGTCTTGGTCGTGGCAAAGGTCGAAAAGGGCAAGTCGAAAAAGCTCAAATCG<br>GTCAAAGAGTTGTTGGGTATTACGATTATGGAGCGGTCCTCGTTCGAAAAGAACCCC<br>ATCGACTTTCTGGAAGCCAAAGGCTATAAGGAAGTCAAGAAGGATCTCATCATCAAA<br>TTGCCGAAATATTCCCTCTTTGAATTGGAAAACGGTCGTAAGCGTATGCTGGCCTCG<br>GCGAAGCAACTGCAGAAAGGGAACGAATTGGCGCTCCCTTCCAAGTATGTGAACTTC<br>CTCTACCTCGCGTCGCACTACGAGAAGTTGAAAGGGTCGCCAGAAGATAATGAACAA<br>AAGCAGCTCTTTGTGGAGCAGCACAAGCATTACCTCGACGAGATTATCGAGCAAATT<br>AGCGAGTTCAGCAAACGGGTCATTCTCGCCGACGCCAATCTCGACAAAGTGCTGTCC<br>GCATACAATAAGCATCGGGACAAACCAATTCGGGAGCAAGCCGAAAATATCATCCAC<br>CTGTTTACCTTGACCCGCCTGGGTGCCCCTCGCGCGTTCAAGTACTTTGATACCACC<br>ATCGACCCAAAACAGTACCGGTCCACGAAGGAGGTCCTCGATGCTACCCTCATCCAT<br>CAGAGCATTACCGGCCTCTATGAGACGCGCATCGACCTCTCG**TGA** |
| Sequence of the *APOBEC* cytidine deaminase | **ATG**AGTAGCGAAACCGGTCCGGTTGCAGTGGATCCGACCCTGCGCCGTCGCATTGAA<br>CCGCACGAGTTTGAAGTGTTCTTTGATCCGCGCGAGCTGCGCAAAGAAACTTGCCTG<br>CTGTACGAGATTAACTGGGGTGGCCGCCATAGCATCTGGCGCCATACCAGCCAGAAC<br>ACCAATAAGCACGTGGAAGTGAATTTTATTGAAAAATTTACCACCGAGCGCTACTTC<br>TGCCCTAATACCCGCTGCAGCATCACCTGGTTTCTGAGCTGGAGCCCGTGCGGCGAA<br>TGTAGTCGCGCCATTACCGAGTTCCTGAGCCGCTATCCGCATGTGACCCTGTTCATC<br>TACATCGCCCGTCTGTACCATCACGCCGATCCGCGCAATCGCCAAGGTCTGCGTGAT<br>CTGATTAGCAGCGGTGTGACCATCCAGATCATGACCGAACAAGAGAGCGGCTATTGC<br>TGGCGCAACTTCGTGAACTATTCTCCGAGCAACGAAGCCCACTGGCCGCGTTATCCG<br>CATCTGTGGGTGCGCCTGTATGTGCTGGAGCTGTACTGCATCATCCTGGGCCTGCCG<br>CCTTGCCTGAATATTCTGCGCCGTAAACAGCCGCAACTGACATTCTTCACCATCGCA<br>CTGCAGAGCTGCCATTATCAGCGCCTGCCGCCGCACATTTTATGGGCCACCGGTCTG<br>AAA**TAA** |
| Sequence of the *ABEmax* module | **ATG**ATGAGCGAAGTCGAGTTCTCCCATGAGTATTGGATGCGGCACGCTCTGACCCTC<br>GCGAAACGTGCTCGCGACGAGCGGGAAGTCCCAGTGGGCGCTGTCCTCGTGCTGAAT<br>AACCGCGTGATTGGCGAAGGCTGGAATCGGGCCATCGGACTCCATGATCCTACGGCA<br>CACGCAGAAATCATGGCCCTGCGCCAGGGTGGCCTCGTCATGCAAAATTATCGGCTG |

| | |
|---|---|
| | TACGACGCCACGTTGTATTCCACCTTCGAGCCCTGCGTCATGTGTGCCGGGGCAATG<br>ATTCACTCCCGTATTGGGCGTGTGGTGTTCGGGGTGCGCAATGCAAAGACCGGCGCT<br>GCCGGCTCGCTCATGGATGTCTTGCACCATCCCGGTATGAACCACCGGGTCGAGATT<br>ACCGAAGGCATTTTGGCCGACGAGTGCGCCGCGCTCTTGTGCCGGTTTTTTCGTATG<br>CCACGGCGTGTGTTCAACGCTCAAAAGAAAGCACAAAGCTCCACCGATTCCGGGGGT<br>TCCTCGGGCGGCTCGAGCGGGTCCGAAACCCCTGGTACGTCGGAATCGGCTACGCCA<br>GAATCGTCCGGCGGCTCGTCCGGCGGTTCT**TAA** |
| Synthetic ABE-editing module (ABEmax-SpRY) | **ATG**AGCGAAGTCGAGTTCTCCCATGAGTATTGGATGCGGCACGCTCTGACCCTCGCG<br>AAACGTGCTCGCGACGAGCGGGAAGTCCCAGTGGGCGCTGTCCTCGTGCTGAATAAC<br>CGCGTGATTGGCGAAGGCTGGAATCGGGCCATCGGACTCCATGATCCTACGGCACAC<br>GCAGAAATCATGGCCCTGCGCCAGGGTGGCCTCGTCATGCAAAATTATCGGCTGTAC<br>GACGCCACGTTGTATTCCACCTTCGAGCCCTGCGTCATGTGTGCCGGGGCAATGATT<br>CACTCCCGTATTGGGCGTGTGGTGTTCGGGGTGCGCAATGCAAAGACCGGCGCTGCC<br>GGCTCGCTCATGGATGTCTTGCACCATCCCGGTATGAACCACCGGGTCGAGATTACC<br>GAAGGCATTTTGGCCGACGAGTGCGCCGCGCTCTTGTGCCGGTTTTTTCGTATGCCA<br>CGGCGTGTGTTCAACGCTCAAAAGAAAGCACAAAGCTCCACCGATTCCGGGGGTTCC<br>TCGGGCGGCTCGAGCGGGTCCGAAACCCCTGGTACGTCGGAATCGGCTACGCCAGAA<br>TCGTCCGGCGGCTCGTCCGGCGGTTCTGACAAGAAGTACAGCATCGGCCTGGCCATC<br>GGCACCAACTCTGTGGGCTGGGCCGTGATCACCGACGAGTACAAGGTGCCCAGCAAG<br>AAATTCAAGGTGCTGGGCAACACCGACCGGCACAGCATCAAGAAGAACCTGATCGGA<br>GCCCTGCTGTTCGACAGCGGCGAAACAGCCGAGAGAACCCGGCTGAAGAGAACCGCC<br>AGAAGAAGATACACCAGACGGAAGAACCGGATCTGCTATCTGCAAGAGATCTTCAGC<br>AACGAGATGGCCAAGGTGGACGACAGCTTCTTCCACAGACTGGAAGAGTCCTTCCTG<br>GTGGAAGAGGATAAGAAGCACGAGCGGCACCCCATCTTCGGCAACATCGTGGACGAG<br>GTGGCCTACCACGAGAAGTACCCCACCATCTACCACCTGAGAAAGAAACTGGTGGAC<br>AGCACCGACAAGGCCGACCTGCGGCTGATCTATCTGGCCCTGGCCCACATGATCAAG<br>TTCCGGGGCCACTTCCTGATCGAGGGCGACCTGAACCCCGACAACAGCGACGTGGAC<br>AAGCTGTTCATCCAGCTGGTGCAGACCTACAACCAGCTGTTCGAGGAAAACCCCATC<br>AACGCCAGCGGCGTGGACGCCAAGGCCATCCTGTCTGCCAGACTGAGCAAGAGCAGA<br>CGGCTGGAAAATCTGATCGCCCAGCTGCCCGGCGAGAAGAAGAATGGCCTGTTCGGA<br>AACCTGATTGCCCTGAGCCTGGGCCTGACCCCCAACTTCAAGAGCAACTTCGACCTG<br>GCCGAGGATGCCAAACTGCAGCTGAGCAAGGACACCTACGACGACGACCTGGACAAC<br>CTGCTGGCCCAGATCGGCGACCAGTACGCCGACCTGTTTCTGGCCGCCAAGAACCTG<br>TCCGACGCCATCCTGCTGAGCGACATCCTGAGAGTGAACACCGAGATCACCAAGGCC<br>CCCCTGAGCGCCTCTATGATCAAGAGATACGACGAGCACCACCAGGACCTGACCCTG<br>CTGAAAGCTCTCGTGCGGCAGCAGCTGCCTGAGAAGTACAAAGAGATTTTCTTCGAC<br>CAGAGCAAGAACGGCTACGCCGGCTACATTGACGGCGGAGCCAGCCAGGAAGAGTTC<br>TACAAGTTCATCAAGCCCATCCTGGAAAAGATGGACGGCACCGAGGAACTGCTCGTG<br>AAGCTGAACAGAGAGGACCTGCTGCGGAAGCAGCGGACCTTCGACAACGGCAGCATC<br>CCCCACCAGATCCACCTGGGAGAGCTGCACGCCATTCTGCGGCGGCAGGAAGATTTT<br>TACCCATTCCTGAAGGACAACCGGGAAAAGATCGAGAAGATCCTGACCTTCCGCATC<br>CCCTACTACGTGGGCCCTCTGGCCAGGGGAAACAGCAGATTCGCCTGGATGACCAGA<br>AAGAGCGAGGAAACCATCACCCCCTGGAACTTCGAGGAAGTGGTGGACAAGGGCGCT<br>TCCGCCCAGAGCTTCATCGAGCGGATGACCAACTTCGATAAGAACCTGCCCAACGAG<br>AAGGTGCTGCCCAAGCACAGCCTGCTGTACGAGTACTTCACCGTGTATAACGAGCTG<br>ACCAAAGTGAAATACGTGACCGAGGGAATGAGAAAGCCCGCCTTCCTGAGCGGCGAG<br>CAGAAAAAGGCCATCGTGGACCTGCTGTTCAAGACCAACCGGAAAGTGACCGTGAAG<br>CAGCTGAAAGAGGACTACTTCAAGAAAATCGAGTGTTTTGATAGTGTTGAAATTTCA<br>GGAGTTGAAGATAGATTTAATGCTTCATTAGGTACCTACCATGATTTGCTAAAAATT<br>ATTAAAGATAAAGATTTTTTGGATAATGAAGAAAATGAAGATATCTTAGAGGATATT<br>GTTTTAACATTGACCTTATTTGAAGATAGGGAGATGATTGAGGAAAGACTTAAAACA<br>TATGCTCACCTCTTTGATGATAAGGTGATGAAACAGCTTAAACGTCGCCGTTATACT<br>GGTTGGGGACGTTTGTCTCGAAAATTGATTAATGGTATTAGGGATAAGCAATCTGGC<br>AAAACAATATTAGATTTTTTGAAATCAGATGGTTTTGCCAATCGCAATTTTATGCAG<br>CTGATCCATGATGATAGTTTGACATTTAAAGAAGACATTCAAAAAGCACAAGTGTCT<br>GGACAAGGCGATAGTTTACATGAACATATTGCAAATTTAGCTGGTAGCCCTGCTATT<br>AAAAAAGGTATTTTACAGACTGTAAAAGTTGTTGATGAATTGGTCAAAGTAATGGGG<br>CGGCATAAGCCAGAAAATATCGTTATTGAAATGGCACGTGAAAATCAGACAACTCAA<br>AAGGGCCAGAAAAATTCGCGAGAGCGTATGAAACGAATCGAAGAAGGTATCAAAGAA<br>TTAGGAAGTCAGATTCTTAAAGAGCATCCTGTTGAAAATACTCAATTGCAAAATGAA<br>AAGCTCTATCTCTATTATCTCCAAAATGGAAGAGACATGTATGTGGACCAAGAATTA |

| | |
|---|---|
| | GATATTAATCGTTTAAGTGATTATGATGTCGATCACATTGTTCCACAAAGTTTCCTT<br>AAAGACGATTCAATAGACAATAAGGTCTTAACGCGTTCTGATAAAAATCGTGGTAAA<br>TCGGATAACGTTCCAAGTGAAGAAGTAGTCAAAAAGATGAAAAACTATTGGAGACAA<br>CTTCTAAACGCCAAGTTAATCACTCAACGTAAGTTTGATAATTTAACGAAAGCTGAA<br>CGTGGAGGTTTGAGTGAACTTGATAAAGCTGGTTTTATCAAACGCCAATTGGTTGAA<br>ACTCGCCAAATCACTAAGCATGTGGCACAAATTTTGGATAGTCGCATGAATACTAAA<br>TACGATGAAAATGATAAACTTATTCGAGAGGTTAAAGTGATTACCTTAAAATCTAAA<br>TTAGTTTCTGACTTCCGAAAAGATTTCCAATTCTATAAAGTACGTGAGATTAACAAT<br>TACCATCATGCCCATGATGCGTATCTAAATGCCGTCGTTGGAACTGCTTTGATTAAG<br>AAATATCCAAAACTTGAATCGGAGTTTGTCTATGGTGATTATAAAGTTTATGATGTT<br>CGTAAAATGATTGCTAAGTCTGAGCAAGAAATAGGCAAAGCAACCGCAAAATATTTC<br>TTTTACTCTAATATCATGAACTTCTTCAAAACAGAAATTACACTTGCAAATGGAGAG<br>ATTCGCAAACGCCCTCTAATCGAAACTAATGGGGAAACTGGAGAAATTGTCTGGGAT<br>AAAGGGCGAGATTTTGCCACAGTGCGCAAAGTATTGTCCATGCCCCAAGTCAATATT<br>GTCAAGAAAACAGAAGTACAGACAGGCGGATTCTCCAAGGAGTCAATCCGGCCCAAG<br>CGGAACAGCGATAAGCTCATCGCGCGGAAAAAAGATTGGGACCCTAAAAAGTATGGG<br>GGGTTCTTGTGGCCGACGGTCGCTTACAGCGTCTTGGTCGTGGCAAAGGTCGAAAAG<br>GGCAAGTCGAAAAAGCTCAAATCGGTCAAAGAGTTGTTGGGTATTACGATTATGGAG<br>CGGTCCTCGTTCGAAAAGAACCCCATCGACTTTCTGGAAGCCAAAGGCTATAAGGAA<br>GTCAAGAAGGATCTCATCATCAAATTGCCGAAATATTCCCTCTTTGAATTGGAAAAC<br>GGTCGTAAGCGTATGCTGGCCTCGGCGAAGCAACTGCAGAAAGGGAACGAATTGGCG<br>CTCCCTTCCAAGTATGTGAACTTCCTCTACCTCGCGTCGCACTACGAGAAGTTGAAA<br>GGGTCGCCAGAAGATAATGAACAAAAGCAGCTCTTTGTGGAGCAGCACAAGCATTAC<br>CTCGACGAGATTATCGAGCAAATTAGCGAGTTCAGCAAACGGGTCATTCTCGCCGAC<br>GCCAATCTCGACAAAGTGCTGTCCGCATACAATAAGCATCGGGACAAACCAATTCGG<br>GAGCAAGCCGAAAATATCATCCACCTGTTTACCTTGACCCGCCTGGGTGCCCCTCGC<br>GCGTTCAAGTACTTTGATACCACCATCGACCCAAAACAGTACCGGTCCACGAAGGAG<br>GTCCTCGATGCTACCCTCATCCATCAGAGCATTACCGGCCTCTATGAGACGCGCATC<br>GACCTCTCGCAGCTGGGGGGTGAT**TGA** |
| Synthetic CBE-editing module (APOBEC1-SpRY-2UGI) | **ATG**AGTAGCGAAACCGGTCCGGTTGCAGTGGATCCGACCCTGCGCCGTCGCATTGAA<br>CCGCACGAGTTTGAAGTGTTCTTTGATCCGCGCGAGCTGCGCAAAGAAACTTGCCTG<br>CTGTACGAGATTAACTGGGGTGGCCGCCATAGCATCTGGCGCCATACCAGCCAGAAC<br>ACCAATAAGCACGTGGAAGTGAATTTTATTGAAAAATTTACCACCGAGCGCTACTTC<br>TGCCCTAATACCCGCTGCAGCATCACCTGGTTTCTGAGCTGGAGCCCGTGCGGCGAA<br>TGTAGTCGCGCCATTACCGAGTTCCTGAGCCGCTATCCGCATGTGACCCTGTTCATC<br>TACATCGCCCGTCTGTACCATCACGCCGATCCGCGCAATCGCCAAGGTCTGCGTGAT<br>CTGATTAGCAGCGGTGTGACCATCCAGATCATGACCGAACAAGAGAGCGGCTATTGC<br>TGGCGCAACTTCGTGAACTATTCTCCGAGCAACGAAGCCCACTGGCCGCGTTATCCG<br>CATCTGTGGGTGCGCCTGTATGTGCTGGAGCTGTACTGCATCATCCTGGGCCTGCCG<br>CCTTGCCTGAATATTCTGCGCCGTAAACAGCCGCAACTGACATTCTTCACCATCGCA<br>CTGCAGAGCTGCCATTATCAGCGCCTGCCGCCGCACATTTTATGGGCCACCGGTCTG<br>AAAAGCGGTAGTGAAACTCCGGGCACAAGCGAAAGCGCAACCCCGGAAAGTGACAAG<br>AAGTACAGCATCGGCCTGGCCATCGGCACCAACTCTGTGGGCTGGGCCGTGATCACC<br>GACGAGTACAAGGTGCCCAGCAAGAAATTCAAGGTGCTGGGCAACACCGACCGGCAC<br>AGCATCAAGAAGAACCTGATCGGAGCCCTGCTGTTCGACAGCGGCGAAACAGCCGAG<br>AGAACCCGGCTGAAGAGAACCGCCAGAAGAAGATACACCAGACGGAAGAACCGGATC<br>TGCTATCTGCAAGAGATCTTCAGCAACGAGATGGCCAAGGTGGACGACAGCTTCTTC<br>CACAGACTGGAAGAGTCCTTCCTGGTGGAAGAGGATAAGAAGCACGAGCGGCACCCC<br>ATCTTCGGCAACATCGTGGACGAGGTGGCCTACCACGAGAAGTACCCCACCATCTAC<br>CACCTGAGAAAGAAACTGGTGGACAGCACCGACAAGGCCGACCTGCGGCTGATCTAT<br>CTGGCCCTGGCCCACATGATCAAGTTCCGGGGCCACTTCCTGATCGAGGGCGACCTG<br>AACCCCGACAACAGCGACGTGGACAAGCTGTTCATCCAGCTGGTGCAGACCTACAAC<br>CAGCTGTTCGAGGAAAACCCCATCAACGCCAGCGGCGTGGACGCCAAGGCCATCCTG<br>TCTGCCAGACTGAGCAAGAGCAGACGGCTGGAAAATCTGATCGCCCAGCTGCCCGGC<br>GAGAAGAAGAATGGCCTGTTCGGAAACCTGATTGCCCTGAGCCTGGGCCTGACCCCC<br>AACTTCAAGAGCAACTTCGACCTGGCCGAGGATGCCAAACTGCAGCTGAGCAAGGAC<br>ACCTACGACGACGACCTGGACAACCTGCTGGCCCAGATCGGCGACCAGTACGCCGAC<br>CTGTTTCTGGCCGCCAAGAACCTGTCCGACGCCATCCTGCTGAGCGACATCCTGAGA<br>GTGAACACCGAGATCACCAAGGCCCCCCTGAGCGCCTCTATGATCAAGAGATACGAC<br>GAGCACCACCAGGACCTGACCCTGCTGAAAGCTCTCGTGCGGCAGCAGCTGCCTGAG<br>AAGTACAAAGAGATTTTCTTCGACCAGAGCAAGAACGGCTACGCCGGCTACATTGAC |

GGCGGAGCCAGCCAGGAAGAGTTCTACAAGTTCATCAAGCCCATCCTGGAAAAGATG
GACGGCACCGAGGAACTGCTCGTGAAGCTGAACAGAGAGGACCTGCTGCGGAAGCAG
CGGACCTTCGACAACGGCAGCATCCCCCACCAGATCCACCTGGGAGAGCTGCACGCC
ATTCTGCGGCGGCAGGAAGATTTTTACCCATTCCTGAAGGACAACCGGGAAAAGATC
GAGAAGATCCTGACCTTCCGCATCCCCTACTACGTGGGCCCTCTGGCCAGGGGAAAC
AGCAGATTCGCCTGGATGACCAGAAAGAGCGAGGAAACCATCACCCCCTGGAACTTC
GAGGAAGTGGTGGACAAGGGCGCTTCCGCCCAGAGCTTCATCGAGCGGATGACCAAC
TTCGATAAGAACCTGCCCAACGAGAAGGTGCTGCCCAAGCACAGCCTGCTGTACGAG
TACTTCACCGTGTATAACGAGCTGACCAAAGTGAAATACGTGACCGAGGGAATGAGA
AAGCCCGCCTTCCTGAGCGGCGAGCAGAAAAAGGCCATCGTGGACCTGCTGTTCAAG
ACCAACCGGAAAGTGACCGTGAAGCAGCTGAAAGAGGACTACTTCAAGAAAATCGAG
TGTTTTGATAGTGTTGAAATTTCAGGAGTTGAAGATAGATTTAATGCTTCATTAGGT
ACCTACCATGATTTGCTAAAAATTATTAAAGATAAAGATTTTTTGGATAATGAAGAA
AATGAAGATATCTTAGAGGATATTGTTTTAACATTGACCTTATTTGAAGATAGGGAG
ATGATTGAGGAAAGACTTAAAACATATGCTCACCTCTTTGATGATAAGGTGATGAAA
CAGCTTAAACGTCGCCGTTATACTGGTTGGGGACGTTTGTCTCGAAAATTGATTAAT
GGTATTAGGGATAAGCAATCTGGCAAAACAATATTAGATTTTTTGAAATCAGATGGT
TTTGCCAATCGCAATTTTATGCAGCTGATCCATGATGATAGTTTGACATTTAAAGAA
GACATTCAAAAAGCACAAGTGTCTGGACAAGGCGATAGTTTACATGAACATATTGCA
AATTTAGCTGGTAGCCCTGCTATTAAAAAAGGTATTTTACAGACTGTAAAAGTTGTT
GATGAATTGGTCAAAGTAATGGGGCGGCATAAGCCAGAAAATATCGTTATTGAAATG
GCACGTGAAAATCAGACAACTCAAAAGGGCCAGAAAAATTCGCGAGAGCGTATGAAA
CGAATCGAAGAAGGTATCAAAGAATTAGGAAGTCAGATTCTTAAAGAGCATCCTGTT
GAAAATACTCAATTGCAAAATGAAAAGCTCTATCTCTATTATCTCCAAAATGGAAGA
GACATGTATGTGGACCAAGAATTAGATATTAATCGTTTAAGTGATTATGATGTCGAT
CACATTGTTCCACAAAGTTTCCTTAAAGACGATTCAATAGACAATAAGGTCTTAACG
CGTTCTGATAAAAATCGTGGTAAATCGGATAACGTTCCAAGTGAAGAAGTAGTCAAA
AAGATGAAAAACTATTGGAGACAACTTCTAAACGCCAAGTTAATCACTCAACGTAAG
TTTGATAATTTAACGAAAGCTGAACGTGGAGGTTTGAGTGAACTTGATAAAGCTGGT
TTTATCAAACGCCAATTGGTTGAAACTCGCCAAATCACTAAGCATGTGGCACAAATT
TTGGATAGTCGCATGAATACTAAATACGATGAAAATGATAAACTTATTCGAGAGGTT
AAAGTGATTACCTTAAAATCTAAATTAGTTTCTGACTTCCGAAAAGATTTCCAATTC
TATAAAGTACGTGAGATTAACAATTACCATCATGCCCATGATGCGTATCTAAATGCC
GTCGTTGGAACTGCTTTGATTAAGAAATATCCAAAACTTGAATCGGAGTTTGTCTAT
GGTGATTATAAAGTTTATGATGTTCGTAAAATGATTGCTAAGTCTGAGCAAGAAATA
GGCAAAGCAACCGCAAAATATTTCTTTTACTCTAATATCATGAACTTCTTCAAAACA
GAAATTACACTTGCAAATGGAGAGATTCGCAAACGCCCTCTAATCGAAACTAATGGG
GAAACTGGAGAAATTGTCTGGGATAAAGGGCGAGATTTTGCCACAGTGCGCAAAGTA
TTGTCCATGCCCCAAGTCAATATTGTCAAGAAAACAGAAGTACAGACAGGCGGATTC
TCCAAGGAGTCAATCCGGCCCAAGCGGAACAGCGATAAGCTCATCGCGCGGAAAAAA
GATTGGGACCCTAAAAAGTATGGGGGGTTCTTGTGGCCGACGGTCGCTTACAGCGTC
TTGGTCGTGGCAAAGGTCGAAAAGGGCAAGTCGAAAAAGCTCAAATCGGTCAAAGAG
TTGTTGGGTATTACGATTATGGAGCGGTCCTCGTTCGAAAAGAACCCCATCGACTTT
CTGGAAGCCAAAGGCTATAAGGAAGTCAAGAAGGATCTCATCATCAAATTGCCGAAA
TATTCCCTCTTTGAATTGGAAAACGGTCGTAAGCGTATGCTGGCCTCGGCGAAGCAA
CTGCAGAAAGGGAACGAATTGGCGCTCCCTTCCAAGTATGTGAACTTCCTCTACCTC
GCGTCGCACTACGAGAAGTTGAAAGGGTCGCCAGAAGATAATGAACAAAAGCAGCTC
TTTGTGGAGCAGCACAAGCATTACCTCGACGAGATTATCGAGCAAATTAGCGAGTTC
AGCAAACGGGTCATTCTCGCCGACGCCAATCTCGACAAAGTGCTGTCCGCATACAAT
AAGCATCGGGACAAACCAATTCGGGAGCAAGCCGAAAATATCATCCACCTGTTTACC
TTGACCCGCCTGGGTGCCCCTCGCGCGTTCAAGTACTTTGATACCACCATCGACCCA
AAACAGTACCGGTCCACGAAGGAGGTCCTCGATGCTACCCTCATCCATCAGAGCATT
ACCGGCCTCTATGAGACGCGCATCGACCTCTCGCAGCTGGGGGGTGATAGCGGCGGG
AGCGGCGGGAGCGGGGGGAGCACTAACCTCAGCGACATTATCGAGAAGGAAACCGGG
AAGCAACTGGTGATTCAGGAGAGCATCCTGATGCTCCCCGAGGAGGTGGAGGAAGTG
ATCGGGAACAAGCCGGAGAGCGACATCTTGGTCCACACCGCCTATGACGAAAGCACC
GATGAGAACGTCATGCTCTTGACGTCCGATGCGCCTGAGTACAAGCCGTGGGCCCTC
GTCATTCAAGACTCCAATGGTGAAAACAAAATTAAGATGCTGAGCGGAGGATCCGGA
GGATCTGGAGGCAGCACTAACCTCAGCGACATTATCGAGAAGGAAACCGGGAAGCAA
CTGGTGATTCAGGAGAGCATCCTGATGCTCCCCGAGGAGGTGGAGGAAGTGATCGGG
AACAAGCCGGAGAGCGACATCTTGGTCCACACCGCCTATGACGAAAGCACCGATGAG

| | |
|---|---|
| | AACGTCATGCTCTTGACGTCCGATGCGCCTGAGTACAAGCCGTGGGCCCTCGTCATT<br>CAAGACTCCAATGGTGAAAACAAAATTAAGATG**TGA** |

**Figure S1. Calibration of a synthetic cytidine base-editor.** Editing was tested both for the target on the genome (Tn*7*::*mCherry*) and on a plasmid (pSEVA2313R). **(A)** Flow cytometry analysis showing population distribution after 24 h of editing when targeting a gene in the chromosome or **(B)** in a plasmid. Editing efficiency (fraction of cells with shifted fluorescence as comparted to a negative control) was estimated based on the population-wide shift in fluorescence signal using the FlowLogicTM software. *a.u.*, arbitrary units.

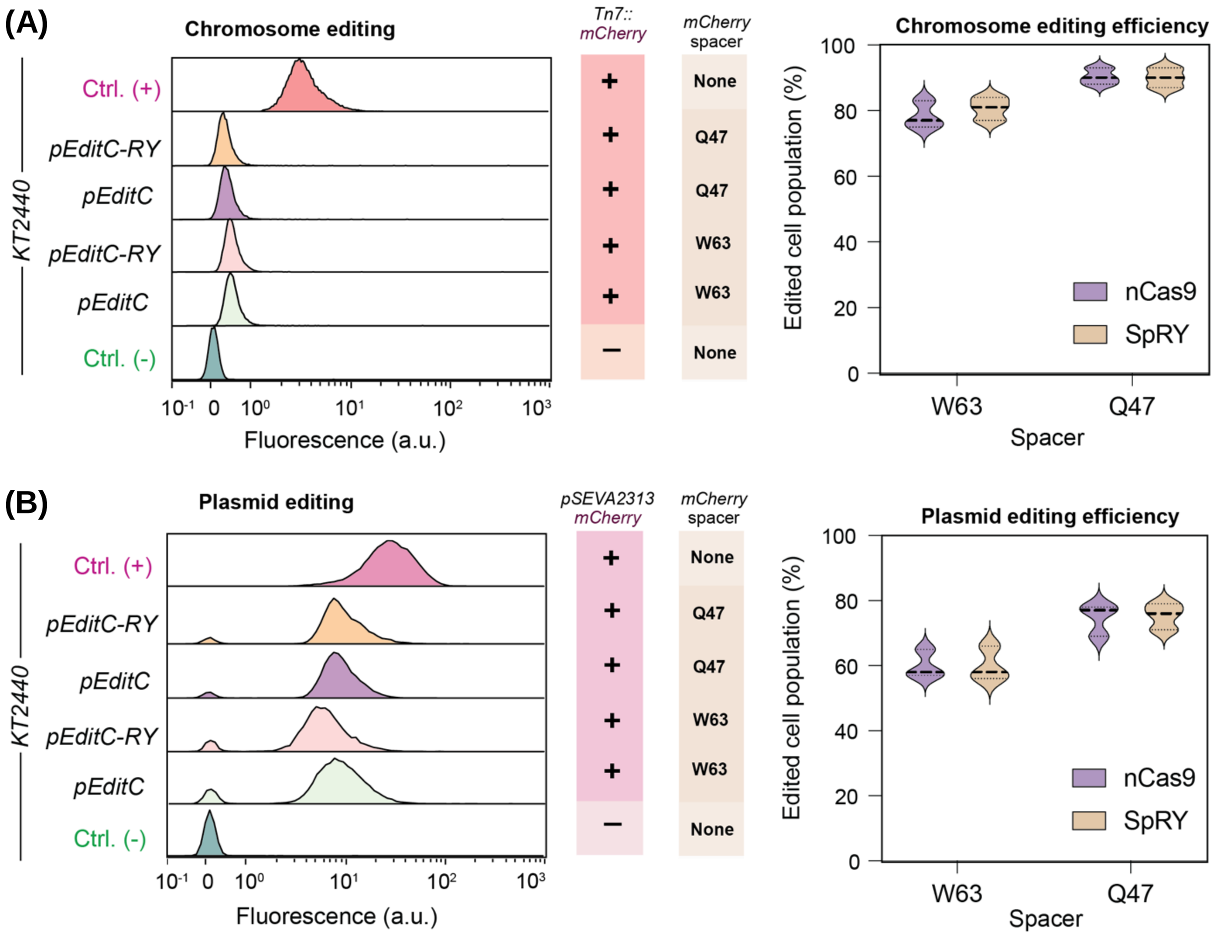

**Figure S2. Calibration of a synthetic adenine base-editor.** Flow cytometry analysis showing the distribution of cell (sub)population(s) after 24 h of editing when targeting a gene in the chromosome to alter the *START* codon with an adenine base-editor. *a.u.*, arbitrary units.

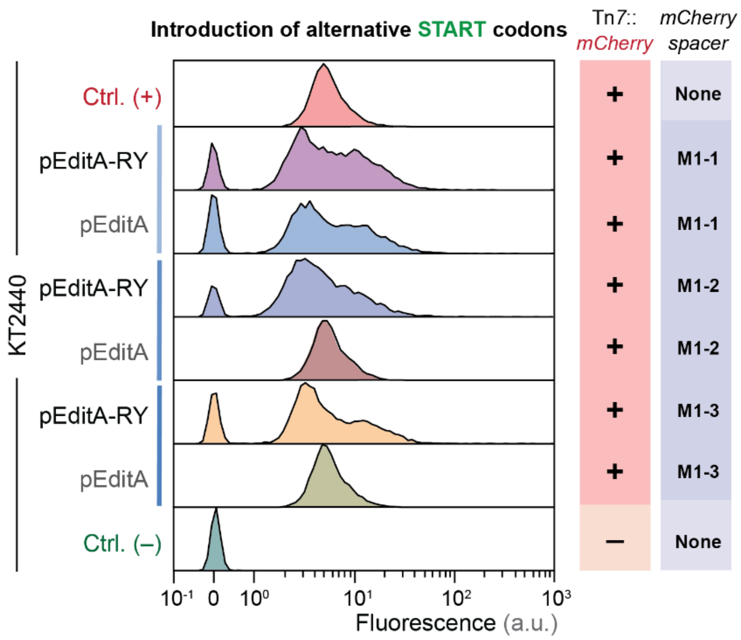

**Figure S3. Frequency of single nucleotide polymorphisms in base-editing experiments.** Samples were taken after 2 rounds of 24-h base-editing of (i) *nicX* with plasmid pCasso·*nicX*$^{Q7}$ (see **Fig. 5C**), followed by vector curing (**CBE_SpRY**); (ii) *mCherry* with plasmid pEditC·*mCherry*$^{W63}$ (see **Fig. 3F**, **CBE_nCas9**); *nicX* gene with plasmid pAblo·*nicX*$^{Q7}$ (see **Fig. 5D**), followed by vector curing (**ABE_SpRY**); and (iv) *mCherry* with plasmid pEditA·*mCherry*$^{M1-1}$ (see **Fig. 3H**, **ABE_nCas9**). *P. putida* KT2440, grown under the same conditions, was used as a control strain and reference genome. Taking the T:A→C:G mutation as an example, this category includes mutations from T→C and A→G. When a T→C mutation appears on either strand of the DNA, the A→G mutation will be found in the same position on the complementary strand. Therefore, the T→C and A→G mutations are classified into a single category. Accordingly, all potential SNP mutations could be classified into six categories; the frequency of each type of mutation is shown below for both cytidine and adenine base-editing experiments. The bars represent the average occurrence of each type of mutation in 10-15 randomly-picked colonies from three independent base editing-experiments ± standard deviation.

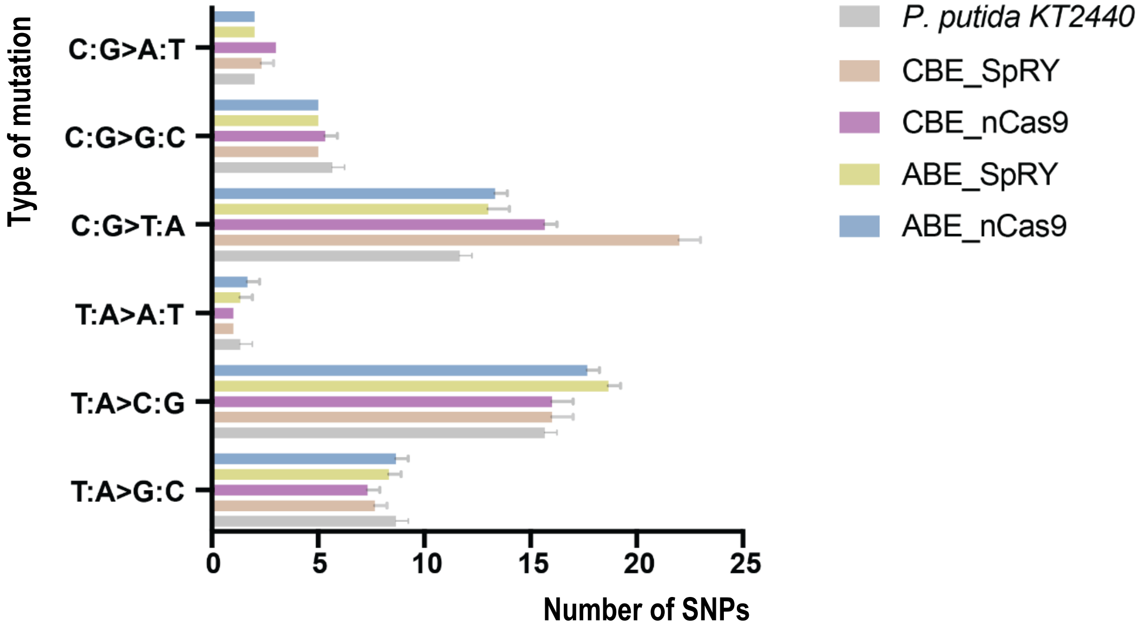
